# Deep mutational scan of a drug efflux pump reveals its structure-function landscape

**DOI:** 10.1101/2021.10.01.462730

**Authors:** Gianmarco Meier, Sujani Thavarasah, Kai Ehrenbolger, Cedric A. J. Hutter, Lea M. Hürlimann, Jonas Barandun, Markus A. Seeger

## Abstract

Drug efflux is a common resistance mechanism found in bacteria and cancer cells. Although several structures of drug efflux pumps are available, they provide only limited functional information on the phenomenon of drug efflux. Here, we performed deep mutational scanning (DMS) on the bacterial ATP binding cassette (ABC) transporter EfrCD to determine the drug efflux activity profile of more than 1500 single variants. These systematic measurements revealed that the introduction of negative charges at different locations within the large substrate binding pocket results in strongly increased efflux activity towards positively charged ethidium, while additional aromatic residues did not display the same effect. Data analysis in the context of an inward-facing cryo-EM structure of EfrCD uncovered a high affinity binding site, which releases bound drugs through a peristaltic transport mechanism as the transporter transits to its outward-facing conformation. Finally, we identified substitutions resulting in rapid Hoechst influx without affecting the efflux activity for ethidium and daunorubicin. Hence, single mutations can convert the ABC exporter EfrCD into a drug-specific ABC importer.

## INTRODUCTION

Multidrug efflux pumps have been studied for more than four decades [1]. Drug efflux is an active transport process requiring the energy of ATP in the case of ABC transporters or ion gradients in the case of secondary-active transporters [2]. A hallmark of drug efflux pumps is their ability to transport a broad range of typically aromatic and cationic compounds. Structural analyses of drug efflux pumps and their regulator proteins revealed that drug binding pockets harbor aromatic amino acids to establish π-π interactions with conjugated drug compounds [3–7]. Yet, in other instances, hydrogen bonds or salt bridges mediated by polar or charged residues were found to be of primary importance for drug recognition [8, 9]. In contrast to the fast progress at the structural front, functional analyses lag behind, because drug transport assays are laborious and thus have to be limited to specific regions of the protein or to certain amino acids used for substitutions, as is the case for alanine scanning [10].

Recent developments in next generation sequencing (NGS) have opened a novel era of protein biochemistry by enabling deep mutational scanning (DMS) analyses. DMS allows studying the effect of hundreds of single variants with regards to protein activity in a single experiment by combining protein libraries with a selection procedure and NGS as a selection read-out [11]. DMS has thus far been mainly used to study protein-protein interactions and catalytic activity profiles of enzymes [12, 13].

Here, we applied DMS to interrogate the sequence-function profile of EfrCD, a heterodimeric type I ABC exporter stemming from the pathogenic bacterium *Enterococcus faecalis.* EfrCD exhibits strong drug efflux activity towards aromatic and cationic drugs such as ethidium, Hoechst 33342 (referred to as Hoechst) and daunorubicin [14]. EfrC and EfrD are expressed as separate half-transporters, which finally heterodimerize to form EfrCD consisting of two transmembrane domains (TMDs) responsible for drug recognition and translocation and two nucleotide binding domains (NBDs), which couple the energy of ATP binding and hydrolysis to conformational changes at the TMDs. EfrCD features asymmetric ATP binding sites. The consensus site is mainly encoded by the EfrD NBD and is competent for ATP hydrolysis. The degenerate site is mainly comprised by the EfrC NBD. Its catalytic residues differ from the consensus sequence and allosterically regulate the consensus site [15, 16]. While the NBDs of EfrCD have been studied extensively [15], it remains elusive how the TMDs recognize and extrude a broad spectrum of drugs out of the cell.

## RESULTS

### Cryo-EM structure of EfrCD and DMS library design

To identify residues lining the transmembrane cavity, we determined the structure of EfrCD using single-particle cryo-EM. The structure of EfrCD was obtained in complex with a nanobody, Nb_EfrCD#1, at an overall resolution of 4.3 Å (Fig. 1a, Fig. S1, Fig. S2, Table S1). EfrCD exhibits a classical type I ABC exporter fold and adopts an inward-facing conformation with the cavity exposed to the cytosol and the NBDs in close proximity to each other. The nanobody binds to the extracellular part of EfrCD and reaches into a crevice formed between EfrC and EfrD, involving mostly hydrophobic interactions contributed by all three CDRs (Fig. S3a). Nb_EfrCD#1 binds with high affinity (K_*D*_ = 7 nM, Fig. S3b) and inhibits ATPase activity of EfrCD by approximately 70 % (Fig. S3c). Based on the cryo-EM structure of EfrCD, we identified 67 residues lining the large inward-facing cavity formed by the TMDs (Fig. 1). To study their role in transporting the EfrCD substrates daunorubicin, ethidium and Hoechst, we mutated them one-by-one to all other 19 amino acids and thereby created a library of 1273 cavity variants. In addition, we mutated 14 highly conserved residues of the asymmetric NBDs of EfrCD, which had been previously studied at the functional level [15]. For these NBD residues, we expected loss-of-function phenotypes for most substitutions regardless of which drug is used in the assay. In summary, our deep mutational scan included a total of 81 residues corresponding to 1539 single variants.

**Figure 1.**
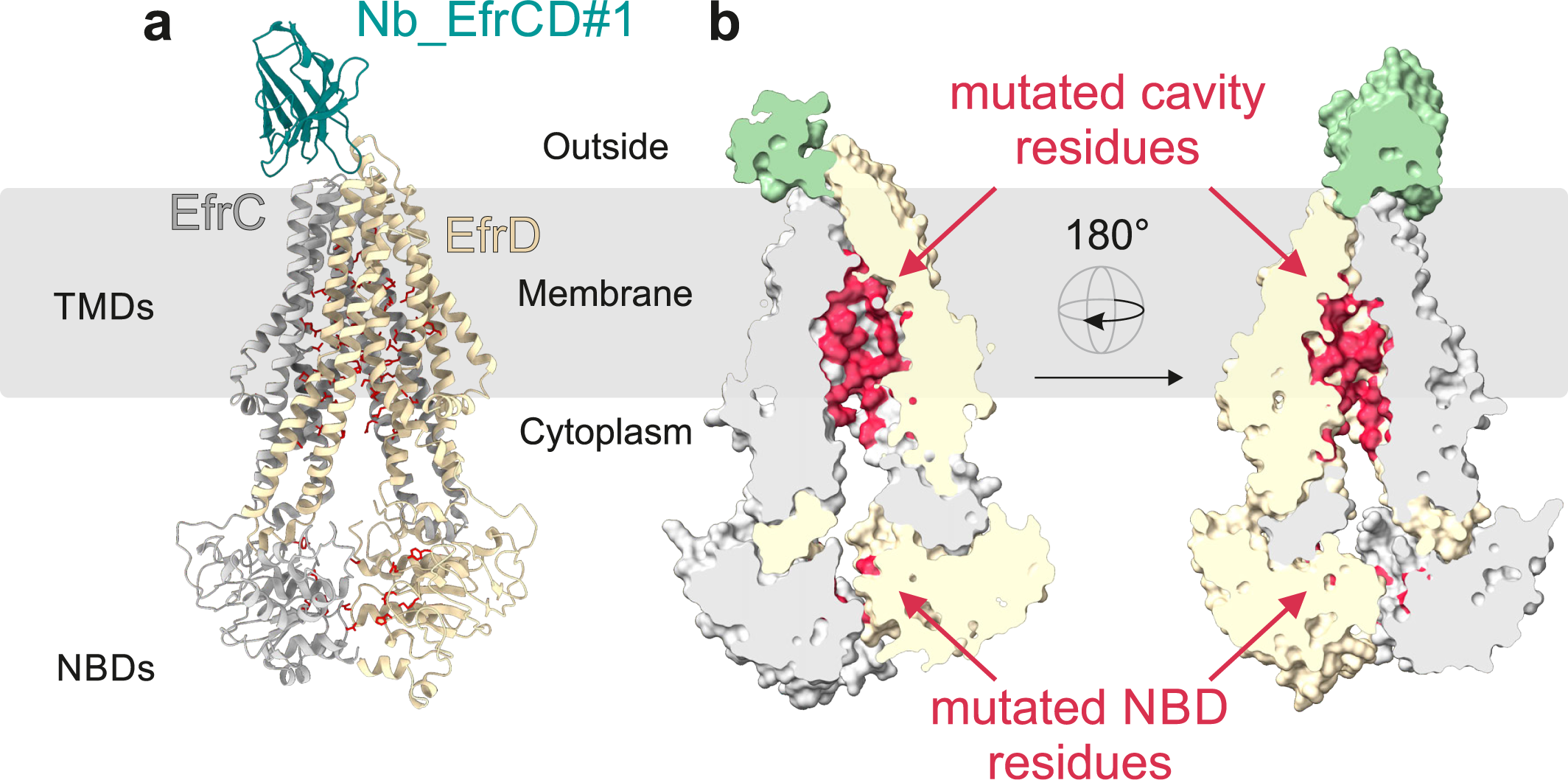
Cryo-EM structure of EfrCD. **a,** Cartoon representation of EfrCD (EfrC in silver and EfrD in light yellow) in complex with nanobody Nb_EfrCD#l (in turquoise). **b,** Cross-sections of EfrCD to visualize the drug binding cavity. Amino acids subjected to DMS are colored in red (67 in the cavity and 14 in the NBDs).

### DMS pipeline overview

To perform DMS experiments, three boundary conditions need to be met. Firstly, an input mutational library has to be generated (Fig. 2a, Fig. S4). As we planned to study substitutions introduced at 81 positions, we had to limit our DMS library to single variants, because a combinatorial library would have been by far too large to be analyzed (Supplementary Note 1). Secondly, a genotype-phenotype linkage is needed to exert selection pressure on each library member. In our case, expression of EfrCD in *Lactococcus lactis* (*L. lactis*) provides resistance towards toxic dyes, resulting in varying growth rates depending on which EfrCD variant is expressed in the individual cell (Fig. S5). The genetic information for the respective EfrCD variant is encoded on a plasmid (Fig. 2b, Supplementary Note 2). Our expression construct adds a GFP C-terminally to EfrD (Fig. 2a) to measure EfrCD production levels in a facile manner. This allows identifying loss-of-function variants that are not produced because of substitutions leading to protein misfolding. Thirdly, a high-throughput read-out is needed to count each variant in the input library as well as after the selection experiment to calculate variant scores (Fig. 2c, Fig. S6). This is achieved by next generation sequencing, in our case Illumina NovaSeq (Supplementary Note 3).

**Figure 2.**
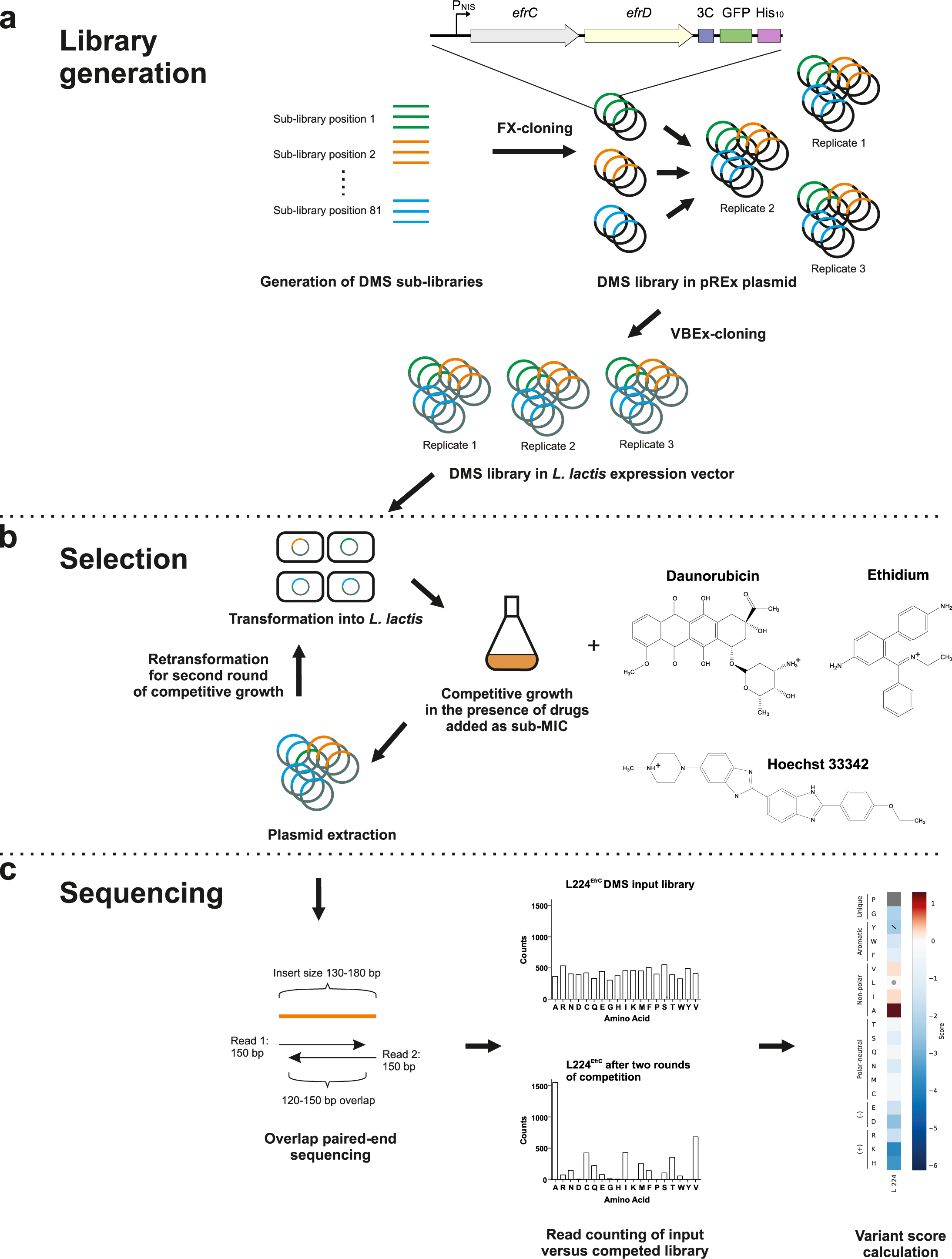
Deep mutational scanning pipeline for drug efflux pump EfrCD. **a,** DMS library preparati on. Three replicates of the DMS library were generated and used in all experiments. **b,** Competitive growth of *L. lactis* expressing EfrCD variants in the presence of three different drugs. **c,** Next generation sequencing of input libraries and competed libraries to determine variant scores for every variant.

### Sequence-function map of EfrCD determined for three drugs

A deep mutational scan of EfrCD was performed in the presence of daunorubicin, Hoechst or ethidium and gave rise to three sequence-function heat maps (Fig. 3). For the majority of the variants (84.4%, 76.6% and 76.7% for the daunorubicin, Hoechst and ethidium, respectively), variant scores were calculated by the program Enrich2 [17]. For around 3% of variants, our NGS data suffered from near-cognate reading errors and the respective scores were not calculated (grey squares in Fig. 3, see also Supplementary note 3). For another 13-20 % of variants, depletion was so pronounced that they were not detected by NGS after two rounds of competition. Consequently, variant scores could not be calculated in these cases of strong depletion (dark blue squares in Fig. 3).

**Figure 3.**
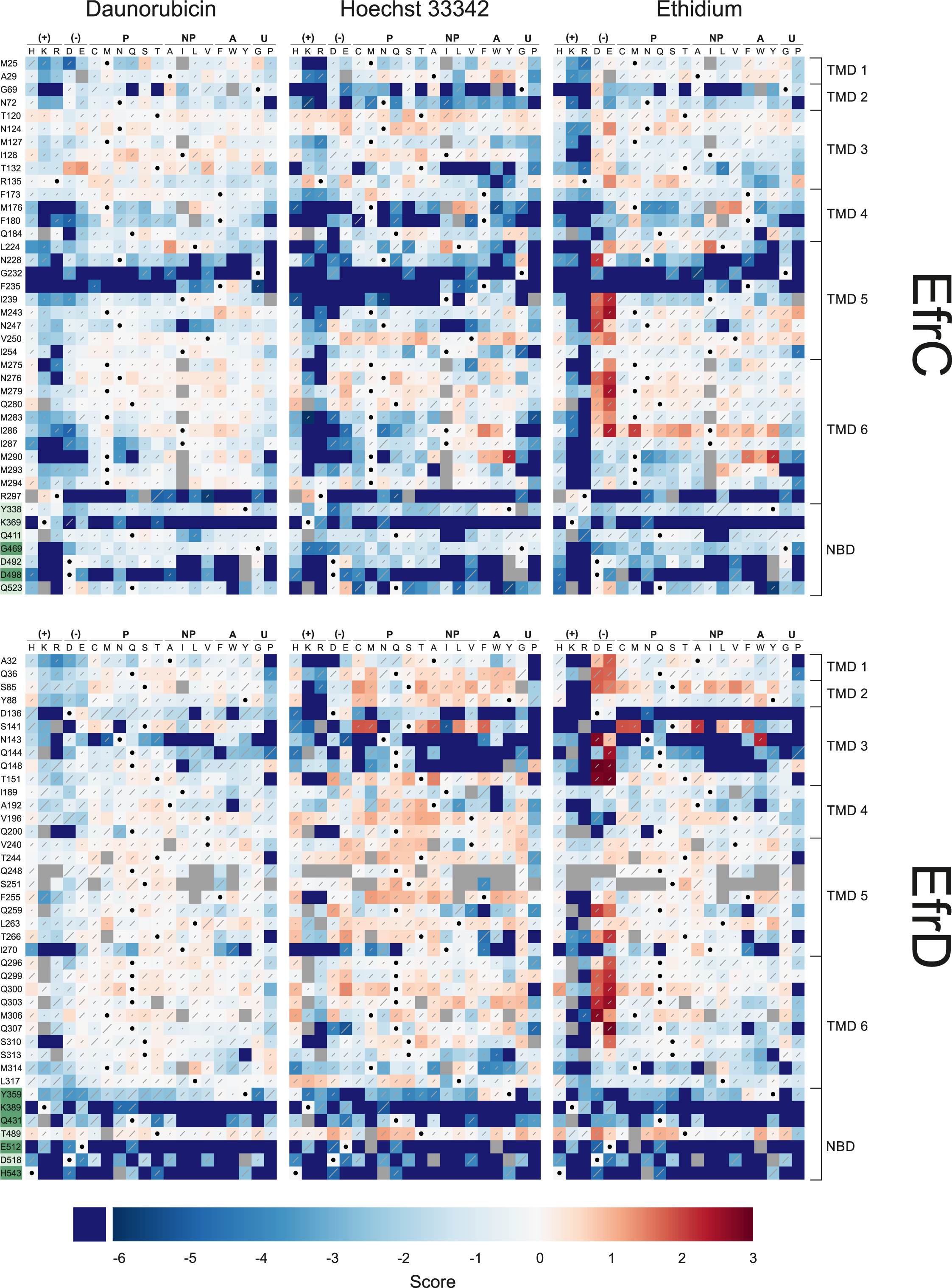
Sequence-function map of EfrCD. Enrich2 software was used to calculate variant scores of the deep mutational scan of EfrCD (EfrC top and EfrD lower panel) performed in the presence of daunorubicin (left), Hoechst (middle) or ethidium (right). Enriched variants are colored in red and depleted variants in blue. Diagonal lines in each cell correspond to the standard error for the variant score and are scaled such that the highest standard error on the plot covers the entire diagonal. Cells with dots have the wild-type amino acid at that position. Dark blue squares represent variants for which no NGS count was detected in at least one of the three competed libraries due to strong depletion. Grey squares denote variants for which no score was calculated either due to high sequencing error frequencies at the specific triplet position, or low input library counts (see also Supplementary note 3). Randomized residues are indicated on the left of the heat map. NBD residues of the consensus nucleotide binding site are shaded in dark green and residues of the degenerate site in light green. Substituting amino acids are indicated on the top and are grouped into positively charged(+), negatively charged(-), polar-neutral (P), non-polar (NP), aromatic (A) and unique (U).

### Validation of DMS approach on the basis of conserved NBD residues

As expected, the great majority of the NBD variants exhibited decreased transport activity (Fig. 4a), including many strongly depleted variants for which no variant score could be determined. Strongly depleted variants are particularly abundant in the EfrD NBD, which harbors most residues of the consensus ATP binding site, whereas they were clearly less frequent in the EfrC NBD containing the degenerate ATP binding site (Fig. 4a). Further, within the NBDs, the sequence-function patterns were highly similar for the three drugs (Fig. 3). In agreement with a previously published study on EfrCD performed with single variants [15], substitutions of the Walker A lysines (K369^EfrC^ and K389^EfrD^), the D-loop aspartates (D498^EfrC^ and D518^EfrD^), the consensus site Walker B glutamate (E512^EfrD^) and the consensus site switch loop histidine (H543^EfrD^) resulted in particularly strong depletion. The A-loop tyrosines (Y338^EfrC^ and Y359^EfrD^), which provide base stacking interactions with the adenine ring of ATP, tolerated aromatic substitutions without losing transport activity (Fig. 3). Again, in agreement with our previous analyses, substitutions are generally well-tolerated in the degenerate site Q-loop (Q411^EfrC^) and in both ABC signature motifs (G469^EfrC^ and T489^EfrD^). Intriguingly, many substitutions at the degenerate site ABC signature motif (T489^EfrD^) were beneficial, as was also shown previously at the example of the T489G^EfrD^ single variant [15]. Hence, the DMS results obtained here are in close agreement with previously investigated single NBD variants of EfrCD, which we consider as a strong indicator for excellent data quality of the entire sequence-function map.

**Figure 4.**
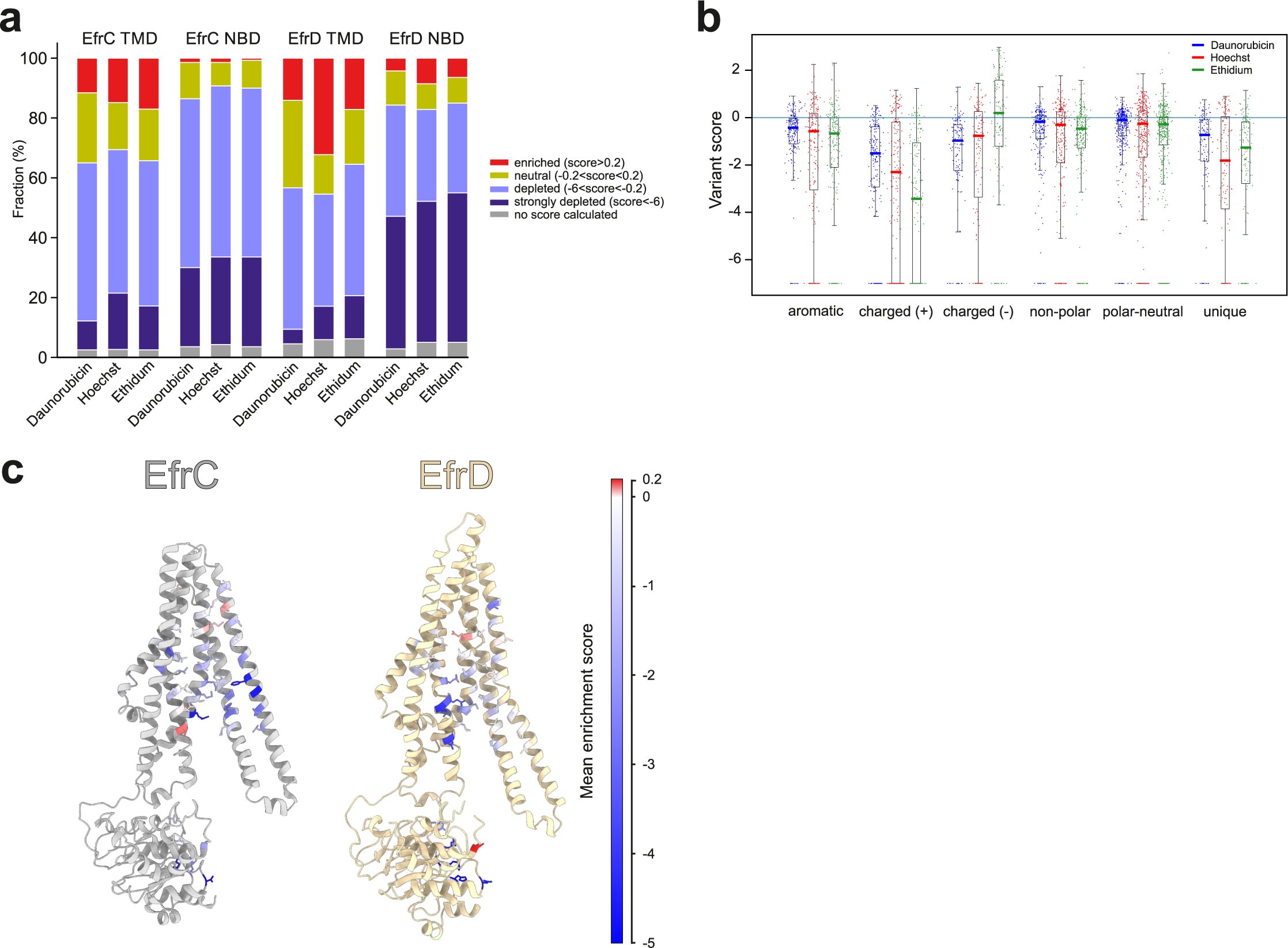
DMS analysis of EfrCD. **a,** Percentage of enriched, neutral, depleted or strongly depleted variants for each domain (EfrC TMD, EfrC NBD, EfrD TMD and EfrD NBD) as determined by DMS in the presence of daunorubicin, Hoechst or ethidium. **b,** Boxplot representationof variant scores (TMD residues only) for daunorubicin (blue), Hoechst (red) and ethidium (green) grouped by properties (aromatic, positively charged, negatively charged, non-polar, polar-neutral and unique) of the substituting amino acid. Shown is the median (coloreline), the interquartile range (boxes) and the last data point in the 1.5-fold interquartile range (whiskers). The score of strongly depleted variants was arbitrarily set to -7. **c,** Mean variant scores of mutated residues (values of all three drugs cumulated) are depicted as colored sticks on the structure of EfrC (left panel in silver) and EfrD (right panel in light yellow). EfrD is turned by 180° in relation to EfrC. The colors of mutated residues correspond to the scale shown onthe right.

### The drug binding cavity can be optimized for drug efflux activity

Substitution of residues in the TMDs lining the drug binding cavity were much better tolerated than in the highly conserved NBDs. As seen in the example of EfrD TMD, up to 30% of substitutions resulted in similar or higher Hoechst efflux activity compared to wild-type EfrCD (score> 0.2) (Fig. 4a). This indicates that the drug efflux activity of wild-type EfrCD can be further increased and that this transporter evolved to transport a still unknown physiological substrate and not drugs and dyes.

### Chemical hallmarks of the drug binding cavity

A high proportion of substitutions to polar-neutral and non-polar amino acids had beneficial effects on transport activity, while introducing positive charges resulted in a comparatively high proportion of inactive variants, in particular with regards to ethidium transport (Fig. 4b). In contrast, the introduction of negative charges into the TMDs had an overall positive impact on ethidium transport (Fig. 3, Fig. 4b) and was also better tolerated with regards to Hoechst and daunorubicin transport than the introduction of positive charges. While all three drugs are positively charged at physiological pH (Fig. 2b), the positive charge of ethidium cannot be removed by deprotonation due to a quaternary nitrogen within the conjugated ring-system [18]. This might explain why in particular for ethidium, electrostatic attraction via the introduction of negatively charged glutamates or aspartates strongly increased drug efflux activity for many variants, while electrostatic repulsion via positively charged residues was poorly tolerated. The residues whose substitutions to negative charges resulted in enrichment upon ethidium exposure primarily locate in the upper half of the inward-facing binding cavity (Fig. S7). Intriguingly, it does not appear to be relevant where exactly the beneficial positive charges are introduced, demonstrating that EfrCD’s drug binding cavity is highly plastic. Next to charge compensation, aromatic residues have been previously shown to play a key role in establishing π-π interactions with conjugated ring systems as they are present in the three drugs used in this study [4, 7]. However, our DMS scan did not reveal an overall beneficial effect upon the introduction of additional aromatic residues into the substrate cavity (Fig. 4b). Nevertheless, some of the aromatic EfrCD variants (M243F^EfrC^, M290Y^EfrC^, L263Y^EfrD^) consistently increased the efflux activity towards all three drugs (Fig. 3), indicating that opposed to the negatively charged residues, the exact location of aromatic residues appears to matter. Wild-type EfrCD contains five aromatic residues in the substrate binding pocket (F173^EfrC^, F180^EfrC^, F235^EfrC^, Y88^EfrD^ and F255^EfrD^), which is a low number when compared to other drug efflux ABC exporters such as ABCB1 [6, 7, 19]. Of these aromatic residues, only two (F180^EfrC^ and F235^EfrC^) appear to be of functional importance, as they did not tolerate substitutions well, with the exception of other aromatic amino acids (Fig. 3). Surprisingly, the introduction of glycines or prolines into the TMDs is often well tolerated and, in some cases, even beneficial and the median variant scores for these unique amino acids were found to be higher than for positively charged residues (Fig. 4b).

### Six residues form a high affinity drug binding site

To identify structure-function relationships, the mean variant scores determined over all substitutions and drugs were visualized in the context of the EfrCD structure (Fig. 4c). This analysis revealed a cluster of six positions (F180^EfrC^, N228^EfrC^, G232^EfrC^, F235^EfrC^, D136^EfrD^ and N143^EfrD^), whose substitution results in a strong depletion for all tested drugs. This region, which we call the “depleted cluster”, is located in the lower half of the substrate binding cavity and involves residues of transmembrane helices (TMHs) 4 and 5 of EfrC and TMH 3 of EfrD (Fig. 5a). TMH 4 and 5 of EfrC constitute the two domain-swapped helices, which cross-over to EfrD and interact with the NBD of EfrD via the coupling helix. Thereby, the consensus nucleotide binding site has the capacity to cross-communicate with the depleted cluster via EfrD’s highly conserved Q-loop. Another important structural aspect of the depleted cluster is the fact that this region becomes partially inaccessible as EfrCD transits from its inward-facing to its outward-facing conformation (Fig. S8a-c). To address the molecular underpinning of the depleted cluster, we generated the corresponding single alanine variants for these six positions by site-directed mutagenesis. In addition, we randomly picked 46 depleted cluster variants from our DMS library (Table S2). In a first analysis, we assessed growth in the presence of daunorubicin and expression levels via GFP fluorescence measurements of all depleted cluster variants. This analysis revealed that independently of the substituting amino acid, the great majority of variants exhibit decreased daunorubicin transport activity while most of them maintained normal expression levels (Table S2). In a second step, we thoroughly analyzed the six alanine variants as well as a subset of seven randomly chosen variants with regards to i) drug resistance in growth assays ii) drug efflux based on Hoechst and ethidium fluorescence and iii) ATPase activity measurements using purified and nanodiscs-reconstituted ErfrCD variants (Fig. 5, Fig. S9, Fig. S10). To increase clarity, we show the results for a subset of the variants exhibiting characteristic phenotype patterns in Fig. 5 and the remaining results in Fig. S9. Of note, growth assays for all six alanine variants and the F235Q^EfrC^ and N143S^EfrD^ variants in the presence of the three drugs were performed as biological triplicates and the complete data is shown in Fig. S10.

**Figure 5.**
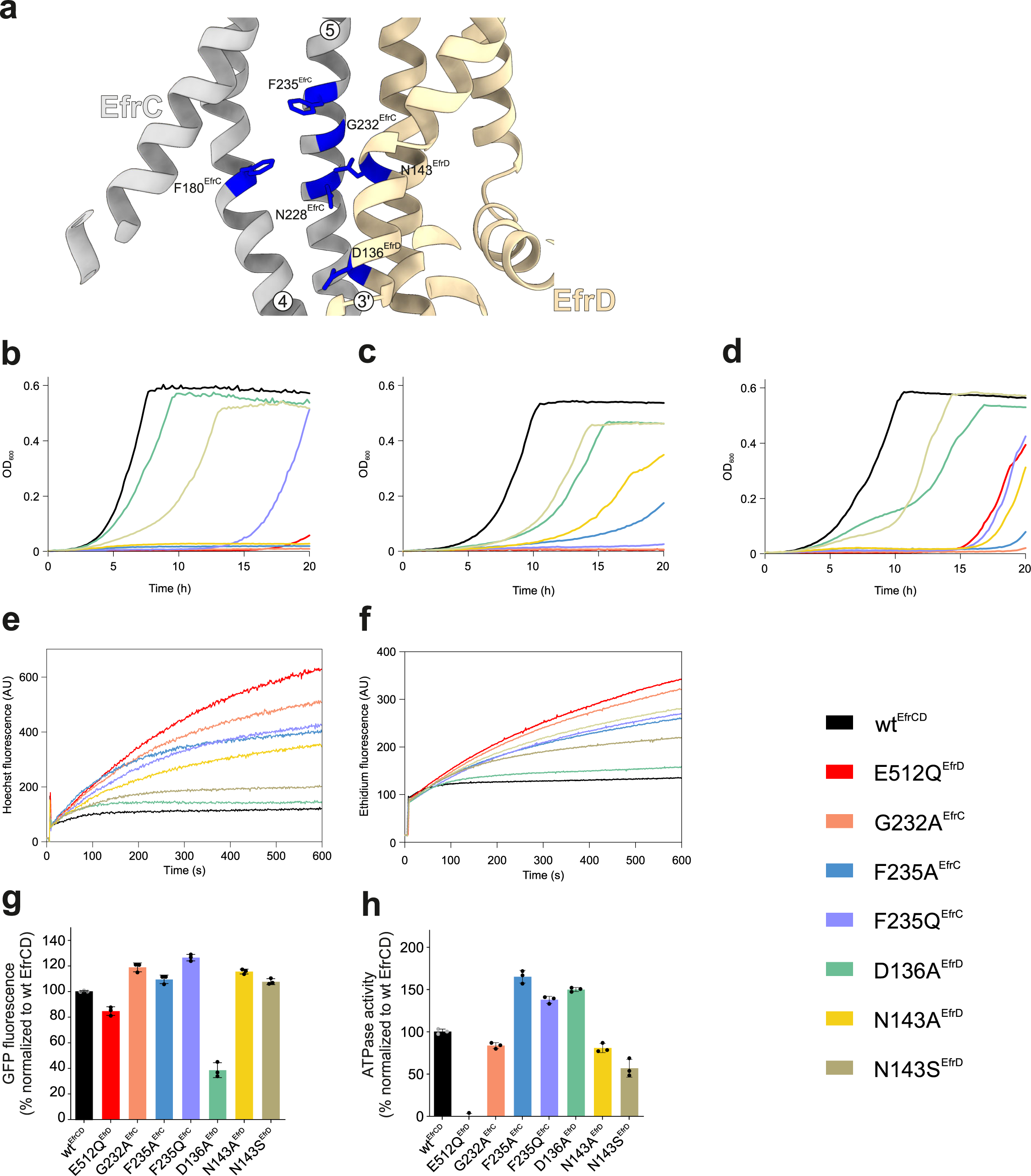
Single clone analysis of depleted cluster variants. **a,** Depleted cluster residues (colored in blue) localize on TMHs 4 and 5 of EfrC (silver) and on TMH 3 of EfrD (light yellow). **b-d,** Growth curves of depleted cluster variants determined in the presence of 8 µM daunorubicin **(b),** 16 µM ethidium **(c)** and 1.5 µM Hoechst **(d). e, f,** Transport assay with fluorescent substrates Hoechst **(e)** and ethidium **(f).** Fluorescence spectroscopy was used to monitor the accumulation of drugs by *L. lactis* expressing the respective EfrCD variant. Drug efflux manifests in slower increase of fluorescence. Representative data of two biological replicates are shown. **g,** Protein production level of depleted cluster variants relative to wild-type EfrCD (100 %) as determined by GFP fluorescence in *L. lactis* expressing the variants fused to GFP. **h,** Basal ATPase activity of purified EfrCD variants measured in nanodiscs. The signal from the inactive E512Q^EfrD^ variant was subtracted and shown is the relative ATPase activity to wild-type EfrCD (100 %). Wild-type EfrCD and the inactive variant E512Q^EfrD^ were included as control in all assays. Error bars correspond to standard deviations of technical triplicates. Growth curves shown in **b-d** are representative data of biological triplicates (all curves shown in Fig. S10). Fluorescence measurements shown in **e** and **f** are representative data of biological duplicates.

In agreement with the DMS results, the ability of all variants to grow in the presence of all drugs was decreased (Fig. 5b-d, Fig. S9a, Fig. S10). Next, we analyzed whether the variants were able to mediate Hoechst and ethidium efflux when overexpressed in *L. lactis*. To this end, we monitored the uptake of these fluorescent dyes in intact *L. lactis* cells. This assay format is highly sensitive in distinguishing residual activities of strongly impaired variants [15] and thus cannot be directly compared with the DMS results generated based on cellular growth in the presence of drugs. We observed variable degrees of activity loss versus wild-type EfrCD, with some variants (G232A^EfrC^, G232D^EfrC^, F235A^EfrC^ and F235Q^EfrC^) exhibiting almost completely abolished efflux activity (Fig. 5e and f, Fig. S9b and c). To quantify protein production levels, we exploited the C-terminal GFP tag as a reporter for expression of EfrCD (Fig. 5g, Fig. S9d). Variants D136A^EfrD^ and D136I^EfrD^ clearly showed diminished EfrCD production, while the other variants had similar production levels as the one observed for wild-type EfrCD. Since *L. lactis* expressing variants D136A^EfrD^ or D136I^EfrD^ still conferred considerable resistance towards drugs (Fig. 5b-d, Fig. S9a), EfrCD produced at a substantially decreased level appears to be sufficient to confer drug resistance. Hence, for the EfrCD variants that were no longer capable of supporting growth in the presence of daunorubicin, insufficient transporter production cannot explain the loss of function. To assess if these substitutions interfere with ATP hydrolysis, the respective variants were overexpressed in *L. lactis* and purified in detergent. In agreement with the GFP-fluorescence data, all eight variants could be purified to homogeneity using size exclusion chromatography (SEC) (Fig. S11). The fraction of the main SEC peak eluting at around 11.4 ml was used for reconstitution into nanodiscs, and basal ATPase activities were determined (Fig. 5h, Fig. S9e). With the exception of variants N143S^EfrD^ and G232D^EfrC^, whose ATPase activity was diminished by around 40% and 80 %, respectively, relative to wild-type EfrCD, the ATPase activities of the remaining six depleted cluster variants were found to be fully intact. Of special mention are variants F180A^EfrC^, F180K^EfrC^, F180T^EfrC^, N228A^EfrC^, N228V^EfrC^, F235A^EfrC^ and F235Q^EfrC^, which were expressed at similar levels as wild-type EfrCD in *L. lactis* (Fig S9d), exhibited equal or slightly higher ATPase activities (Fig. 9e), but were clearly impaired with regards to drug efflux (Fig. 3). Our analyses thus suggest that the residues of the depleted cluster directly interact with the transported drugs and thereby define a high affinity binding site for drugs when EfrCD adopts its inward-facing state.

### Specificity determining clusters

In a search for specificity-determining residues, we screened the DMS data for EfrCD variants exhibiting a low variant score (<-3) for one drug and a high variant score (>-0.2) for at least one other drug (Fig. 6a-c). Substitutions leading to a pronounced decrease of ethidium efflux (ethidium specificity variants) were observed in a cluster of seven residues (N124R/K^EfrC^, M294R^EfrC^, S141E^EfrD^, Q144R/T^EfrD^, V196R^EfrD^, Q307F^EfrD^ and L317R^EfrD^), which are located in the lower part of the TMDs and form a belt around the cavity (Fig. 6d). Many of these variants introduce positive charges and presumably result in electrostatic repulsion with ethidium’s positive charge at the entry of EfrCD’s drug binding cavity. In contrast, positive charges at this cavity entry belt are well tolerated with regards to daunorubicin or Hoechst transport, presumably because the positive charge in these molecules can be removed by transient deprotonation and is not located and partially delocalized in the conjugated ring systems as is the case for ethidium (Fig. 2b). Single clone analyses of five of these ethidium specificity variants (N124R^EfrC^, M294R^EfrC^, S141E^EfrD^, Q144T^EfrD^ and Q307F^EfrD^) confirmed the DMS data (Fig. S12).

**Figure 6.**
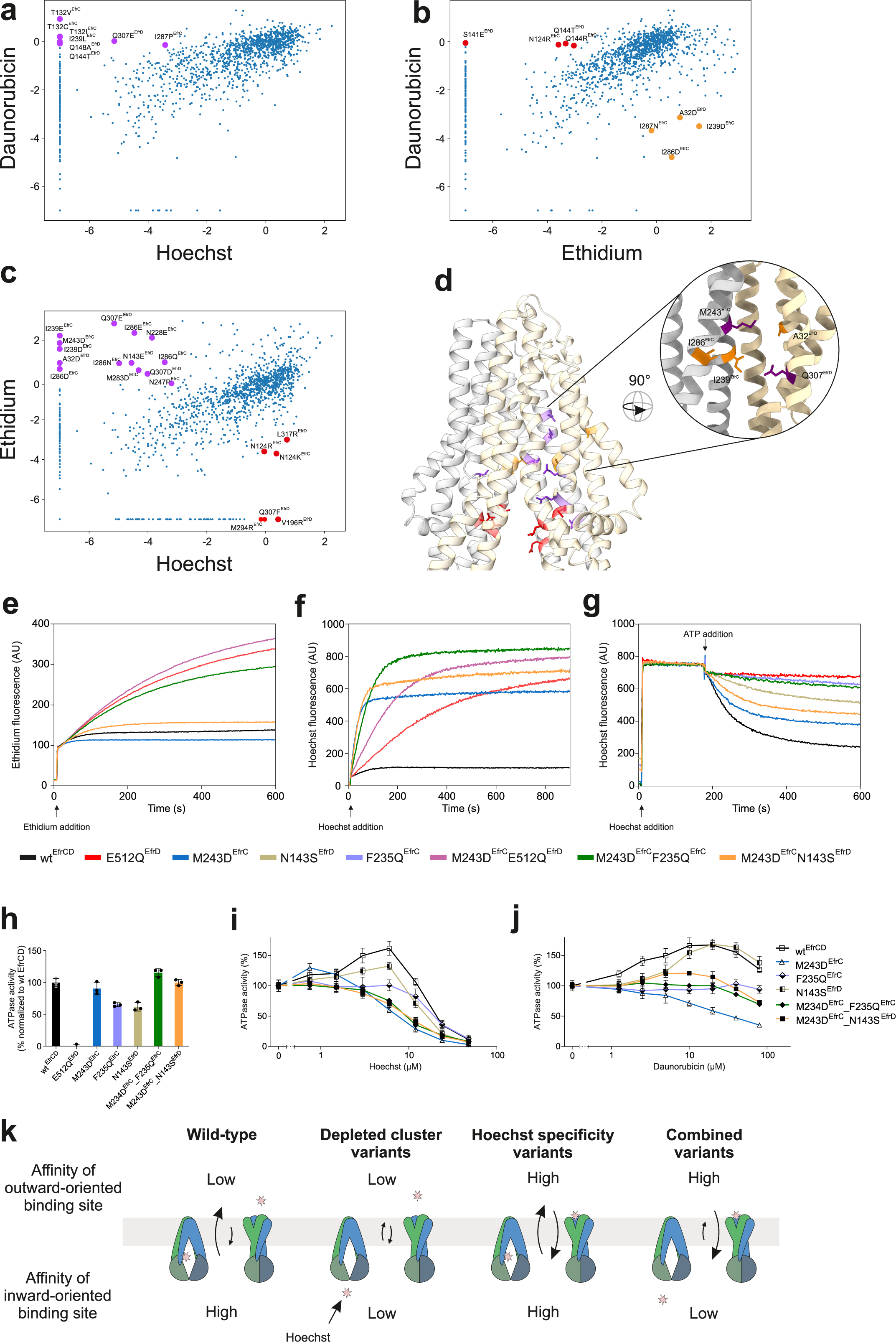
Specificity variants investigated at a single clone level. a, b, c, Variant scores for daunorubicin versus Hoechst **(a),** daunorubicin versus ethidium **(b)** and Hoechst versus ethidium **(c).** Transporter variants depleted (score <-3) for one drug but neutral or enriched (score >-0.2) for at least one of the other drugs are categorized as specificity variants. Specificity variants for daunorubicin (orange), Hoechst (purple) and ethidium (red) are shown as colored dots and are labeled with the respective substitution. **d,** Specificity variants are depicted as sticks in the context of the EfrCD structure and are colored in orange (daunorubicin and Hoechst), purple (Hoechst only) or red (ethidium). The close-up view shows the core of the Hoechst/daunorubicin specificity cluster including residues I239^Efrc^, M243^EfrC^, I286^EfrC^, A32^EfrD^ and Q307^EfrD^ **e, f, g,** Fluorescence transport assays. Hoechst **(e)** and ethidium **(f)** accumulation in intact *L. lactis* cells or Hoechst accumulation in ISOVs **(g)** containing overexpressed Hoechst-specificityvariant M243Dmc alone or in combination with depleted cluster variants F235^EfrC^ or N143S^EfrD^. **h, i, j,** Basal ATPase activity **(h)** and drug-modulated ATPase activity **(i** and j) of nanodisc-reconstituted EfrCD variants analyzed in **(e-g).** ATPase activities were measured in the presence of increasing concentrations of Hoechst **(i)** or daunorubicin **(j).** Data were normalized to the basal activity of the respective variant in the absence of drugs. Error bars are standard deviations of technical triplicates. **k,** Schematic interpretationof transport data. For detailed explanations, see main text.

For daunorubicin, four variants met the specificity filtering criteria (I239D^EfrC^, I286D^EfrC^, I287N^EfrC^ and A32D^EfrD^) (Fig. 6a and b). These four variants were also strongly depleted for Hoechst, but not for ethidium. Finally, we identified another nine variants, which lost their capacity to confer resistance to Hoechst, but not to daunorubicin and ethidium (Fig. 6a and c). Hence, there is a large cluster of 13 residues where certain (but not all) substitutions lead to a loss of Hoechst resistance, which overlaps with four residues whose substitutions to mostly aspartates result in daunorubicin sensitivity (Fig. 6d).

### Introduction of a secondary Hoechst binding site results in EfrCD-mediated influx

Within the Hoechst/daunorubicin specificity cluster, a set of five residues (inset of Fig. 6d) caught our attention. Their substitution with negatively charged residues (I239D^EfrC^, M243D^EfrC^, I286D^EfrC^, A32D^EfrD^ and Q307D/E^EfrD^) led to a particularly strong loss of Hoechst and in part also daunorubicin resistance, but were neutral or even beneficial to confer ethidium resistance according to the DMS analysis (Fig. 3). To study these variants in more detail, we generated them individually by site-directed mutagenesis to validate their drug-specific phenotypes at a single clone level in growth assays (Fig. S13a) and drug transport assays with ethidium and Hoechst. In the *in vivo* fluorescence transport assay, these five variants displayed an unexpectedly steep fluorescence increase upon Hoechst addition, which was clearly faster than Hoechst uptake mediated by *L. lactis* cells expressing the ATPase-deficient E512Q^EfrD^ control variant (Fig. 6f, Fig. S13b). This effect was Hoechst-specific, because the same set of variants behaved very similarly to wild-type EfrCD when monitoring ethidium uptake (Fig. 6e, Fig. S13b). Interestingly, these variants did not accumulate the same amount of Hoechst as E512Q^EfrD^. Instead, they reached a steady-state level, akin to the curves obtained with cells expressing wild-type EfrCD, but at a higher intracellular Hoechst concentration. This suggests EfrCD-mediated import of Hoechst at the beginning of the uptake experiment when intracellular drug concentration is low. Once the intracellular concentration reaches a certain level, influx and efflux are in equilibrium with no net transport across the membrane. We propose that in these variants, a secondary high affinity binding site for Hoechst is created, which is exposed when the transporter is in the outward-facing state (Fig. S8d-f). The fact that the most pronounced Hoechst-specificity variants encompassed substitutions with negatively charged residues further lends support to this hypothesis.

We next asked the question, whether ATP-mediated conformational cycling at the NBDs is required for the fast uptake of Hoechst in these variants. To this end, we combined the Hoechst-specificity variant M243D^EfrC^ with the ATPase-deficient E512Q^EfrD^ variant. The respective M243D^EfrC^_E512Q^EfrD^ double variant indeed exhibited a slower initial uptake rate as compared to the M243D^EfrC^ single variant, showing that EfrCD-mediated Hoechst uptake is facilitated by conformational cycling of the transporter (Fig. 6f). In contrast to the M243D^EfrC^ single variant, a steady-state Hoechst level was not reached for the double variant, meaning that influx dominates over efflux during the entire transport experiment, as it is the case for the single E512Q^EfrD^ variant.

### Biochemical characterization of drug binding clusters

Our analyses revealed a drug binding cluster, which is well accessible from the inward-facing cavity and is important for the efflux of all three investigated drugs (depleted cluster). In addition, a secondary Hoechst binding site was found, which is created upon the introduction of mainly negative charges to a cluster accessible in both the inward- and the outward-facing state (Fig. S8d-f). To further investigate the interplay of these two binding sites, we combined the Hoechst-specificity variant M243D^EfrC^ with depleted cluster variants F235Q^EfrC^ or N143S^EfrD^. Performing the Hoechst uptake assay in intact cells, these double variants retained the steep initial increase in fluorescence as it was observed for the M243D^EfrC^ single variant (Fig. 6f). In contrast, the steady-state Hoechst level was higher for both double variants than the M243D^EfrC^ single variant. This can be explained by the diminished capacity of the depleted cluster single variants F235Q^EfrC^ and N143S^EfrD^ to efflux Hoechst (Fig. 5e).

Next, we conducted a Hoechst accumulation assay in inside out vesicles (ISOVs) derived from *L. lactis* cells overexpressing the respective EfrCD variants. In this assay, the ISOVs are first incubated with Hoechst until a stable fluorescent signal is reached due to intercalation of Hoechst into the lipid bilayer, followed by the addition of ATP-Mg, which energizes EfrCD transporters that are facing with their NBDs to the outside. In case of active transport, Hoechst accumulates in the vesicle lumen, which in turn leads to a decrease of fluorescent signal due to the concomitant acidification of the vesicle interior through the proton-translocating F_1_F_0_-ATPase and the protonation of Hoechst inside the vesicle lumen (Fig. S14) [20]. Hoechst transport mediated by the M243D^EfrC^ variant was only mildly impaired relative to ISOVs containing wild-type EfrCD (Fig. 6g). This finding agrees with the notion that this variant is vulnerable to fast Hoechst influx (Fig. 5c), but is still capable of Hoechst efflux, which corresponds to Hoechst pumping into the ISOV in this particular assay (Fig. S14). In contrast, the depleted cluster variant F235Q^EfrC^ was almost completely inactive in terms of Hoechst efflux, whereas the depleted cluster variant N143S^EfrD^ had some residual transport activity (Fig. 6g), which is in line with transporter assays conducted in intact cells (Fig. 5c). The M243D^EfrC^_F235Q^EfrC^ double variant has no transport activity in ISOVs, meaning that the depleted cluster variant F235Q^EfrC^ dominates in this transport assay format. Finally, the M243D^EfrC^_N143S^EfrD^ double variant exhibited an intermediate transport function as compared to the corresponding single variants, suggesting that in this assay format, the Hoechst specificity variant improves the activity of the depleted cluster variant (Fig. 6g). To further gain insights into the underlining mechanism, we purified these variants and reconstituted them into lipid nanodiscs to investigate the drug-induced modulation of ATPase activity. It was thoroughly documented that modulation of the ATPase activity of ABC transporters by substrates is a clear sign of its interaction with the protein [21, 22]. When ATPase activity is plotted against drug concentration, often a bell-shaped curve is observed. The prevailing explanation for this phenomenon is that the initial stimulation of ATPase activity at lower drug concentrations is due to the interaction of the substrate with a high affinity binding site, while the inhibition of ATPase activity at higher substrate concentrations is due to the existence of secondary low affinity substrate binding sites within the transporter [23, 24]. To investigate drug-stimulation of ATPase activity, the EfrCD variants were purified and reconstituted in nanodiscs and exhibited basal ATPase activities similar to wild-type EfrCD (Fig. 6h). In agreement with previous work [14], wild-type EfrCD displayed such a bell-shaped ATPase modulation curve for Hoechst and daunorubicin (Fig. 6i and j). The depleted cluster variants F235Q^EfrC^ and N143S^EfrD^ exhibited a weaker stimulation signal for Hoechst (Fig. 6h) or needed a higher drug concentration to reach maximal stimulation for daunorubicin (Fig. 6i), reinforcing our hypothesis that these variants exhibit diminished affinity at the primary drug binding site.

In contrast, the M243D^EfrC^ variant displayed its stimulation maximum at a Hoechst concentration much lower than in the case of wild-type EfrCD, most likely due to ATPase inhibition at low Hoechst concentrations via the secondary Hoechst binding site introduced by this substitution (Fig. 6i). In the presence of daunorubicin, the ATPase activity of the M243D^EfrC^ variant can no longer be activated, indicating that inhibition at the secondary binding site dominates in this case (Fig. 6j).

As summarized in Fig. 6k, our thorough biochemical analysis of single and double variants revealed the existence of an inward-oriented high affinity drug binding site and an outward-oriented secondary Hoechst binding site that is formed upon introduction of negatively charged residues.

## DISCUSSION

Mutational studies have a long history in the field of ABC transporters. Of particular mention are the systematic mutational analyses of ABCB1 by Loo and Clarke, a project that took more than a decade to study the effect of cysteine mutations introduced at the TMDs [25]. Random mutagenesis and functional screening had also been applied multiple times to characterize drug efflux pumps or ABC transporters. In these studies, yeast ABC transporter Pdr5 mutants with altered drug specificity were identified [26], the *Yarrowia lipolytica* pheromone ABC transporter Ste6 was trained to export the initially poorly exported **a**-factor of *Saccharomyces cerevisiae* [27] and the RND drug efflux pump AcrB was screened for mutants that resisted inhibition by an efflux pump inhibitor [28]. In a recent study, the eleven tryptophane residues of murine ABCB1 were screened for functionally-neutral substitutions using site-saturation mutagenesis to generate a tryptophane-free functional ABCB1 variant for drug interaction studies [29]. Common to these studies is that with the exception of Swartz et al, the mutations were introduced by random mutagenesis (i.e. were untargeted). Further, the screens operated based on positive selection, i.e. only functionally neutral or gain-of-function variants were selected and further analyzed, while loss-of-function mutations were ignored during analysis. The experimental read-out was Sanger sequencing of isolated “top” clones that excelled according to the imposed positive selection criteria and in most cases, the identified variants contained multiple substitutions which needed to be disentangled in follow-up experiments. Without any doubt, these pioneering studies provided the ground for this work, as they demonstrated the evolutionary flexibility of drug efflux pumps and ABC transporters to be taught new functions and to alter substrate specificity.

However, the randomness of the above approaches together with the vast sequence space owing to the combination of mutations prevented a systematic recording of sequence-function maps of these transporters. Our DMS approach overcomes these limitations as it permits for the in-depth characterization of a relatively small, but near-complete set of single-site substitutions. Thereby, novel insights into the sequence-function landscape of the TMDs in terms of drug recognition and drug specificity were gained. We surmise that many of the identified patterns and especially the Hoechst-specificity cluster would have escaped discovery, if a systematic alanine-scan or a random mutagenesis approach had been chosen to analyze EfrCD. A hallmark of DMS is the quantification of both gain- and loss-of-function substitutions under exactly identical conditions. We consider the side-by-side comparison of variants in a single culture flask as a key element to overcome the inherent day-by-day variability of drug transport experiments (as is evident from the three biological replicates of growth curves shown in Fig. S10, Fig. S12 and Fig. S13).

The systematic nature of the DMS approach allowed pinpointing general biophysical features of drug efflux pump substrate cavities. At least in EfrCD, electrostatic interactions can be established via a large number of positions, as is prominently seen for ethidium efflux that benefits from the introduction of negative charges into the upper half of the inward-facing drug binding cavity. In turn, a belt at the cavity entry barely tolerates positive charges in the context of ethidium transport. Considering that π-π interaction mediated by aromatic residues had been reported to be of particular importance to confer polyspecificity to drug efflux pumps [4, 6, 7, 30], we were surprised to observe that the introduction of aromatic residues into the drug binding cavity only occasionally resulted in improved drug efflux.

In a remarkable study on Pdr5, an alanine mutation in the H-loop of the consensus ATP binding site altered the drug specificity of this fungal multidrug transporter [31]. In our DMS data on EfrCD, however, we did not find any substitutions within the NBDs that would alter substrate specificity. Another extensively discussed topic in the field is the asymmetry of heterodimeric ABC transporters, in particular such ones with degenerate ATP binding sites [16, 32, 33]. We clearly noted differences between substitutions at the EfrC and the EfrD chain with regards to the NBDs, which were less well tolerated at residues of EfrD. This was expected, because EfrD residues mainly contribute to the consensus ATP binding site. In contrast, variant scores did not greatly differ between EfrC and EfrD at the level of the TMDs (Fig. 4a). This indicates that residues of both EfrC and EfrD contribute to drug recognition and transport and no notable asymmetries appear to exist at the level of the TMDs.

The DMS analysis revealed a polyspecific high-affinity drug binding site, which is only fully accessible when EfrCD adopts its inward-facing state, but is deformed and more narrow in EfrCD’s outward-facing conformation (Fig. S8a-c). This suggests drug extrusion akin to a peristaltic pump, a mechanism which had been previously suggested for the RND drug efflux pump AcrB [34] and ABCB1 [6]. The systematic DMS approach also revealed a vulnerability of EfrCD, namely a cluster of residues in the upper part of the cavity whose substitution into negatively charged residues results in transporter variants that confer rapid Hoechst influx, but still confer resistance towards ethidium. This cluster likely binds Hoechst with high affinity when EfrCD adopts its outward-facing state and thereby counteracts Hoechst extrusion mediated by the peristaltic pumping via the inward-oriented drug binding site (Fig. 6k). Hence, we discovered a delicate interplay between influx and efflux that finally decides over the ultimate direction of transport. Intriguingly, type I ABC exporters, which mediate siderophore or solute import had been described recently [22, 35]. Further, ABCB1 containing 14 alanine substitutions in TMHs 6 and 12 was found to exhibit active drug import, but lost its capacity to efflux drugs [19]. The work presented here provides insights into the plasticity of the large substrate binding cavity of ABC transporters and shows how even single substitutions can influence transport directionality.

Additional work will be needed to explore a larger number of residues of EfrCD or to compare different drug efflux pumps. Further, it would be attractive in future studies to implement fluorescence-activated cell sorting (FACS) into our DMS workflow to gain information on the transporter expression level in a high-throughput manner [36].

This work provides the basis to explore structure-function relationships of drug efflux pumps. We anticipate that DMS will be widely applied to generate high-content functional data urgently needed to interpret the increasing number of cryo-EM structures of drug efflux pumps at the molecular level.

## MATERIALS AND METHODS

### Cloning

Vector pREXdmsC3GH was generated on the basis of pREXC3GH (Addgene #47077) by introducing inverted SapI sites, which allow to excise the *efrCD* operon by restriction digest using SapI. Using primers pREXC3GH_SapI_for (5’-tat ata GCA GGA AGA GCA TTA GAA GTT TTG TTT CAA GGT CCA CAA TTC) and pREXC3GH_SapI_rev (5’-tat ata ACT AGA AGA GCC CAT GGT GAG TGC CTC CTT ATA ATT TAT TTT GTA G) the vector backbone was PCR-amplified and the resulting fragment was ligated with the ccdB kill-cassette excised from pINIT_cat (Addgene #46858) with SapI. Vector pREXNH3CA used to clone EfrCD in frame with a C-terminal Avi-tag was constructed from pREXNH3 (Addgene #47079) by PCR amplification with 5’ phosphorylated primers pREXNH3(newAvi_5’P)_FW (5’-aga aaa tcg aat ggc acg aaT AAT AAC TAG AGA GCT CAA GCT TTC TTT GA) and pREXNH3(newAvi_5’P)_RV (5’-gag ctt cga aga tat cgt tca gac cTG CAG AAG AGC TGA ACT AGT GG) which insert the Avi-tag in front of the stop codon in pREXNH3. The resulting PCR product was circularized using blunt end ligation. Using FX cloning and SapI restriction digest [37], wild-type or mutant *efrCD* was first sub-cloned into pREXdmsC3GH or pREXNH3CA and from there via vector-backbone exchange (VBEx) using SfiI restriction digest into the nisin-inducible *L. lactis* backbone pERL to finally obtain *L. lactis* expression vectors pNZdmsC3GH or pNZNH3CA, respectively [38]. Single clone variants were introduced by QuikChange mutagenesis on the *efrCD*-containing pREXdmsC3GH plasmid using the primers listed in Table S3 and were verified by Sanger sequencing.

### DMS library preparation

The single site saturation libraries were synthesized by Twist Bioscience using the most abundant *Lactococcus lactis* codons (G:GGT, E:GAA, V:GTT, A:GCT, R:AGA, S:AGT, K:AAA, N:AAT, M:ATG, I:ATT, T:ACA, W:TGG, Y:TAT, L:TTA, F:TTT, C:TGC, Q:CAA, H:CAT, P:CCA, D:GAT) and overhangs containing SapI sites. Libraries were amplified by PCR using Phusion® HF polymerase (Thermo Fisher Scientific) and page-purified primers DMS_EfrCD_for_2 (5’-ACC ATG GGC TCT TCT AGT GAC CTT ATT ATT C) and DMS_EfrCD_rev_2 (5’-TTC TAA TGC TCT TCC TGC TTC AAA AAC AAA TTG ATT TT). A sub-library per randomized position was generated by FX cloning [37] of these PCR products into vector pREXdmsC3GH (see above) using chemically-competent MC1061 cells. After recovery, transformed cells were grown overnight in LB medium containing 0.5% glucose and 100 μg/ml ampicillin. Transformation efficiency was monitored by plating 1:100 of the recovery product onto LB agar plates containing 120 μg/ml ampicillin. Plasmid sub-libraries in the pREXdmsC3GH vector were extracted (QIAprep Spin Miniprep Kit, Qiagen). DNA concentration was determined based on A260 using a nanodrop. Sub-libraries with randomized positions in the *efrC* and *efrD* gene were mixed separately and three replicates of each randomized half-transporter were created. In a first attempt, sub-libraries were mixed equimolarly and the input library was analysed by NGS. This quality check revealed that sub-libraries still contained a variable fraction of wild-type *efrCD*, which resulted in an uneven distribution of variants over all positions (Fig. S4b). To overcome this problem, sub-library ratios were adjusted according to this initial NGS analysis by mixing the sub-libraries (in pREXdmsC3GH vector) in respective ratios. The 81 sub-libraries were mixed in three independent pipetting reaction (three replicates), which were then sub-cloned using VBEx into the *L. lactis* expression vector pNZdmsC3GH [38]. To this end, freshly prepared electro-competent *L.lactis* NZ9000 *ΔlmrCD ΔlmrA* were transformed with VBEx product, recovered and then grown in liquid M17 culture containing 0.5% glucose and 5 μg/ml chloramphenicol. Plating of small aliquots after recovery on M17 agar plates supplemented with 0.5% glucose and 5 μg/ml chloramphenicol revealed that at least 10^6^ colony forming units were obtained per replicate. Thereby, the DMS library size of 1539 variants was oversampled by more than 600-fold to prevent diversity bottlenecks. After overnight growth, glycerol stocks of the three DMS library replicates were prepared as 1 ml aliquots and stored at -80°C.

### Competitive growth

For competition experiments, 50 ml of M17, 0.5% glucose with 5 μg/ml chloramphenicol was inoculated with a glycerol stock (1 ml aliquot) of *L. lactis* containing the DMS library and grown without shaking overnight. 50 ml of fresh M17, 0.5% glucose with 5μg/ml chloramphenicol was inoculated with 500 μl of the overnight culture harbouring the DMS library plus 10 μl overnight culture of wild-type *efrCD* cloned in pNZdmsC3GH. Cells were grown for 2h at 30°C without shaking. Protein expression was induced for 30 min by the addition of a nisin-containing culture supernatant of *L. lactis* NZ9700 added at a dilution of 1:10’000 (v/v). OD_600_ was determined and normalized to 0.5. 24 ml of M17, 0.5% glucose, 5 μg/ml chloramphenicol, containing nisin (1:10’000 (v/v)) with the respective drugs was inoculated with 1:100 induced *L. lactis* cells containing the DMS libraries (OD_600_ of 0.005 at the start of the competition). The following drug concentrations were used: 1.5 µM Hoechst 33342, 16 µM ethidium and 8 µM daunorubicin. Competitive growth was performed at 30 °C without shaking overnight. The next morning, an OD_600_ of 2.5 was measured, meaning that, in average, the cells have divided around 9 times. *L. lactis* cell pellets were collected by centrifugation at 4’000g for 10 min.

### Plasmid extraction

The competed DMS libraries (encoded on the *L. lactis* plasmid pNZdmsC3GH) were extracted by resuspending the cell pellets in 200 μl of 20% sucrose, 10 mM Tris/HCl pH 8.0, 10 mM K-EDTA pH 8.0, 50 mM NaCl followed by addition of 400 μl of 200 mM NaOH, 1% SDS. After inverting the tube 4 times the solution was neutralized by addition of 300 μl of 3 M K-acetate, 5 M acetic acid, pH 4.8. After centrifugation at 20’000g for 10 min, the supernatant was added to 350 μl of EtOH and stored at -20°C for 30 min. The precipitate was centrifuged at 4°C, 20’000g for 5 min and the pellet was washed with 70% EtOH. The pellet was dried at room temperature, resuspended in nuclease-free water containing 20 μg/ml RNaseA (Sigma) and incubated for 30 min at room temperature. The plasmids were purified again using the NucleoSpin Gel and PCR Clean-up Kit (Macherey-Nagel). For the second round of competitive growth, the pNZdmsC3GH plasmids containing the DMS library were transformed into freshly prepared electrocompetent *L. lactis* NZ9000 *ΔlmrCD ΔlmrA* cells and the procedure of competitive growth followed by plasmid extraction was repeated as described above. To extract the *efrCD* genes (encoding the DMS variants) for NGS, 7 μg pNZdmsC3GH plasmid was digested with 40 U of SfiI (NEB), separated on a 1% agarose gel and, the band containing the *efrCD* genes were extracted with the NucleoSpin Gel and PCR Clean-up Kit (Macherey-Nagel).

### Library preparation and NGS using Illumina NovaSeq 6000

500 ng DNA was sheared twice in a Covaris microTUBE on a Covaris E220 Focused-ultrasonicator to achieve a distribution with an average fragment length of 150 base pairs using peak power 175, duty factor 10, cycles/burst 200 for 280s at 7°C. In-between cycles the microTUBEs were spun down before second shearing. 150 ng of sheared fragments was used as input for the TruSeq Nano DNA Library Prep Kit (Illumina). Illumina libraries were prepared following the manufacturer’s protocol with one modification to yield a narrow fragment distribution optimized for 150 bp inserts by using a ratio of 5.4 parts of sample purification beads to one part of water (135 μl SPB + 25 μl nuclease-free H2O) in the library size selection step. Libraries were sequenced on a NovaSeq 6000 system (Illumina) using S1 flow cells and paired-end 2 × 151 cycles sequencing.

### Data analysis

In a first step, adapter sequences, which are a result of the short insert size, were removed using Cutadapt. Adapter-trimmed reads were then aligned to the wild-type EfrCD DNA sequence using the Burrows-Wheeler Aligner [39]. With a home-made Python program, the aligned sequences were then further filtered for sequences with a Phred quality score higher than 30. Reads with insertions or deletions were removed and only overlapping sequences were allowed in the further processing. Variants were counted by allowing only reads with matching sequences in the overlapping region and with a single amino acid substitution. Variant scores were calculated using the software package Enrich2 [17].

### Analysis of single clone variants in growth assays

To determine growth curves of individual EfrCD variants, an overnight culture of *L. lactis* NZ9000 *ΔlmrCD ΔlmrA* cells harbouring mutants in the pNZdmsC3GH vector was used to inoculate 10 ml preculture of M17 0.5% glucose with 5 μg/ml chloramphenicol. Cells were grown for 2 h at 30°C and then protein expression was induced by addition of a nisin-containing culture supernatant of *L. lactis* NZ9700 for 30 min (1:10’000 (v/v)). Cultures were normalized to OD_600_ of 0.5 and used to inoculate 150 μl of fresh medium (1:100 (v/v), M17, 0.5% glucose, 5 μg/ml chloramphenicol containing nisin (1:10’000 v/v)) in 96-well plates. OD_600_ was monitored every 10 min over at least 16 h in a microplate reader at 30°C with shaking between reads.

### Determination of EfrCD production levels via GFP fluorescence

Induced cultures were prepared as described in the growth assay. 2 ml of fresh M17, 0.5% glucose, 5 μg/ml chloramphenicol containing nisin (1:10’000 (v/v)) was inoculated with induced cultures (1:100) and grown overnight at 30°C. Cells were collected by centrifugation at 4’000g for 10 min and washed twice with PBS. Pellets were resuspended in PBS. OD_600_ as well as GFP fluorescence (485 nm excitation, 528 nm emission) was measured in a microplate reader (Cytation 5 BioTek) and the fluorescence signal was normalized to OD_600_. After background subtraction of cells producing EfrCD without GFP tag (autofluorescence of *L. lactis*), expression levels relative to wild-type EfrCD were calculated.

### Fluorescence transport assay

To determine transport activity, *L. lactis* NZ9000 *ΔlmrCD ΔlmrA* cells harbouring EfrCD variants in vector pNZdmsC3GH were grown overnight in M17, 0.5% glucose containing 5 μg/ml chloramphenicol. 40 ml of fresh medium was inoculated with overnight culture (1:50) and grown for 2 h at 30°C to OD_600_=0.4-0.6. Protein expression was initialized by addition of nisin-containing culture supernatant of *L. lactis* NZ9700 (1:1’000 (v/v)) and proceeded for 2 h at 30°C. Cells were collected by centrifugation (4’000g, 10 min, 4°C), washed twice and resuspended in 4 ml of 50 mM K-phosphate pH 7.0, 5 mM MgSO4. Transport assays were carried out exactly as described [14].

### Purification of EfrCD

EfrCD wild-type or variants were expressed in *L. lactis* NZ9000 *ΔlmrA ΔlmrCD* containing plasmid pNZNH3GS, pNZNH3CA or pNZdmsC3GH. Cells were grown in M17, 0.5% glucose, 5 μg/ml chloramphenicol at 30°C to an OD_600_ of 1 and expression was induced with a nisin-containing culture supernatant of *L. lactis* NZ9700 (1:5’000 v/v) for 4h. Membranes were prepared by passing cells four times through a microfluidizer at 35 kpsi in PBS (pH 7.4) containing 15 mM K-EDTA (pH 7.4). After low spin centrifugation (10 min, 8’000g), 30 mM MgCl2 was added to the supernatant and the lysate was incubated with DNase for 30 min at 4°C. After high spin centrifugation (170’000g, 1 h) the pellet was resuspended in TBS (20 mM Tris-HCl pH 7.5, 150 mM NaCl) supplemented with 10% glycerol. Proteins were solubilized with 1% (w/v) n-dodecyl-β-D-maltoside (β-DDM) for 2 h at 4°C. Insolubilized fraction was removed by high spin centrifugation (170’000g, 1 h). The supernatant was supplemented with 30 mM imidazole (pH 7.5) and loaded onto Ni-NTA columns with 2 ml bed volume. When EfrCD was prepared for cryo-EM analyses, nanodisc reconstitutions and ATPase activity assays, detergent was exchanged to n-decyl-β-D-maltoside (β-DM) by washing with 15 bed volumes of 50 mM imidazole pH 7.5, 200 mM NaCl, 10% glycerol and 0.3% (w/v) β-DM. Protein was eluted with 200 mM imidazole pH 7.5, 200 mM NaCl, 10% glycerol and 0.3% (w/v) β-DM and for purifications from pNZNH3GS or pNZdmsC3GH buffer was exchanged to TBS with 0.3% (w/v) β-DM using PD10 columns followed by overnight incubation with 3C protease. When EfrCD was prepared for alpaca immunizations or nanobody selections and analyses, the entire purification was carried out in 0.03% (w/v) β-DDM. For the biotinylated versions expressed from vector pNZNH3CA, the protein was concentrated to 360 µl using Amicon Ultra-4 concentrator units with 50 kDa MWCO. Biotinylation and 3C cleavage was preformed overnight at 4°C in a total reaction volume of 4 ml with 0.2 mg 3C protease and 330 nM BirA (0.016 mg/ml) in buffer containing 20 mM imidazole pH 7.5, 10 mM magnesium acetate, 200 mM ATP, 200 mM NaCl, 10% glycerol, 0.03% (w/v) β-DDM and a 1.2-fold molar excess of biotin.

For all constructs, cleaved His-tag and 3C protease were removed by reverse IMAC and EfrCD was polished by size exclusion chromatography (SEC) on a Superdex 200 Increase 10/300 GL column in 20 mM Tris-HCl pH 7.5, 150 mM NaCl, and 0.3% (w/v) β-DM or 0.03% β-DDM (w/v).

### Nanodisc reconstitution

Membrane scaffold protein MSP1E3D1 was expressed and purified as described [40]. Purified EfrCD proteins were reconstituted into nanodiscs at an EfrCD:MSP:lipid molar ratio of 1:14:500 in Na-HEPES pH 8.0 supplemented with 30 mM cholate. The reconstitution mixture was incubated at 4°C for 30 min. 200 mg Bio-Beads™ SM-2 Resin (Bio-Rad) was added and incubated at 4°C overnight while shaking at 650 rpm. After removal of Bio-Beads™ SM-2 Resin, the reconstituted transporter was further purified by SEC on a Superdex-200 10/300 GL column equilibrated with 20 mM Tris-HCl pH 7.5, 150 mM NaCl.

### ATPase activity assay

ATPase activity was measured using nanodisc-reconstituted EfrCD at a concentration of 2 nM in 20 mM Tris-HCl pH 7.5, 150 mM NaCl, 10 mM MgSO4. For ATPase stimulation, Hoechst 33342 (at final concentrations ranging from 0.75 μM - 48 μM), or daunorubicin (at final concentrations ranging from 1.25 μM - 80 μM) were included. ATPase activities were measured at 30°C for 15 min in the presence of 1 mM ATP and liberated phosphate was detected colorimetrically using the molybdate/malachite green method. 90 μl of the reaction solution was mixed with 150 µl of filtered malachite green solution (10.5 mg/mL ammonium molybdate, 0.5 M H_2_SO_4_, 0.34 mg/mL malachite green and 0.1% Triton X-100). Absorption was measured at 650 nm.

### Nanobody selection and production

Nanobody Nb_EfrCD#1 was generated by immunizing an alpaca with four subcutaneous injections of 200 µg EfrCD in 20 mM Tris-HCl pH 7.5, 150 mM NaCl, and 0.03% (w/v) β-DDM in two-week intervals. Immunizations of alpacas were approved by the Cantonal Veterinary Office in Zurich, Switzerland (animal experiment licence nr. 188/2011). Phages libraries were generated as described previously [41, 42]. Two rounds of phage display were performed on biotinylated EfrCD solubilized with 0.03% (w/v) β-DDM. After the last round of phage display, a 340-fold enrichment was determined by qPCR using AcrB as a negative control. The enriched library was then sub-cloned into the pSb_init vector by FX cloning and 94 clones were screened with ELISA using biotinylated EfrCD as target. Out of 83 positive hits, we found three distinct families based on the CDR composition. Nb_EfrCD#1 was cloned into expression plasmid pBXNPHM3 (Addgene #110099) using FX cloning and expressed as described previously [42]. Tag-free Nb_EfrCD#1 was used in cryo-EM structure determination.

### Affinity determination by Grating-coupled interferometry (GCI)

The affinity of Nb_EfrCD#1 was determined with GCI on the WAVEsystem (Creoptix AG, Switzerland). Avi-tagged EfrCD was captured on a streptavidin PCP-STA WAVEchip (polycarboxylate quasi-planar surface; Creoptix AG) to a density of 2000 pg/mm^2^. For binding kinetics, Nb_EfrCD#1 in 20 mM Tris-HCl pH 7.5, 150 mM NaCl, and 0.03% (w/v) β-DDM was injected with increasing concentrations (0.333, 1, 3, 9, 27 nM) using 50 µl/min flow rate for 120 s at 25°C and dissociation was proceeded for 600 s. Data were analysed on the WAVEcontrol (Creoptix AG) using double-referencing by subtracting the signals from blank injections and from the reference channel and fitted using a Langmuir 1:1 model.

### Cryo-EM structure determination

The cryo-EM sample was prepared by purifying EfrCD, reconstituted in micelles, on a Superose 6 10/300 Increase GL size exclusion column (GE Healthcare) in 20 mM Tris-HCl pH 7.5, 150 mM NaCl 0.15% β-DM at 4°C. Fractions from the monodisperse EfrCD peak were pooled, concentrated to 8 mg/ml, and supplemented with 5 mM MgCl2, 5 mM ATPγS, and Nb_EfrCD#1 in a 1:1 ratio to EfrCD. The supplemented sample was incubated for 20 min on ice prior to grid freezing. Quantifoil R2/1 Cu 200 grids were glow-discharged for 30 s at 50 mA before sample application and flash-freezing in a liquid ethane/propane mix using an FEI Vitrobot Mark IV (Thermo Fisher Scientific), set to a blot force of -5, waiting time of 1 s, and blot time of 4.5 s at 100% humidity and 4°C. Data were collected in three sessions on a Titan Krios (TFS) operated at 300 kV, equipped with a Gatan K2 BioQuantum direct electron detector at the Umeå Core Facility for Electron Microscopy. A total of 4’344 movies, each consisting of 40 frames, were collected with a dose of 52.1 e/Å^2^, a defocus range between -1.5 to -3.3 μm, and a pixel size of 1.04 Å, using the EPU software (Thermo Fisher Scientific) for automated data collection. Initial movie alignment, drift correction, and dose-weighting were done with MotionCor2 [43]. CTFFind-4.1.13 [44] was used for contrast transfer function (CTF) determination. After manual inspection of the micrographs for drift or poor CTF-fits, 3’135 micrographs remained. A total of 720’893 particles, picked using crYOLO [45], were extracted with a box size of 196 pixels (203.84 Å). A 30 Å lowpass-filtered initial model, obtained using cisTEM [46], was used as a reference in a 3D classification with 4 classes using RELION-3.1 [47]. Particles from the top class (35.1%, 254’682 particles) were re-extracted with a larger box size of 296 pixels (307.84 Å) and refined to a resolution of 5.50 Å. CTF-refinement, Bayesian polishing, and consensus 3D refinement resulted in a map of 4.73 Å. The particle pool was exported into cryoSPARC v3.2 [48] and classified into three classes. Those classes were further subjected to heterogeneous refinement resulting in a top class with 212’083 particles. After non-uniform refinement [49] the final map reached an overall resolution of 4.25 Å (Fig. S2).

The resulting EM-density map allowed for model building of the TMD using the high-resolution X-ray structure of the ABC transporter TM287/288 (PDB 4Q4H, 2.53 Å) as a template. The resolution in the peripheral regions (NBD, nanobody) allowed for the placement of homology models. The resulting structure was refined using PHENIX real_space_refine [50] against the final map at 4.25 Å resolution (Table S1) and the side-chain atoms in the less-well resolved peripheral regions (NBDs and nanobody) were removed from the final model. The directional FSC determination was performed using 3DFSC [51], and model validation was performed according to [52]. For model validation, all atomic coordinates were randomly displaced by 0.5 Å, followed by refinement against half map 1. The FSC coefficients of this refined model, and half map 1 or half map 2, were calculated using EMAN2 [53]. Model statistics are presented in Table S1.

### Map and model visualization

Structure analysis and figure preparation were performed using PyMOL (Schrödinger) [54] and UCSF ChimeraX [55].

## DATA AVAILABILITY

The cryo-EM map and coordinates for the EfrCD structure have been deposited in the EM Data Bank with accession code EMD-12816 and the Protein Data Bank with accession code PDB-7OCY. NGS datasets are available on request.

## CODE AVAILABILITY

The code of the Python script used to analyse NGS data was deposited on https://github.com/giameier/DMS_ABC.

## AUTHOR CONTRIBUTIONS

GM and MAS conceived the project. GM generated the DMS libraries, established the selection protocol, programmed and validated the NGS data analysis pipeline and performed the ATPase activity assays. ST generated the great majority of the single clone variants, and analyzed them in growth assays and fluorescence transport assays together with GM. Nanobodies were generated by GM and LH and analyzed by GM and ST. Cryo-EM analyses were performed by KE under the supervision of JB with EfrCD protein purified by CAJH. Cryo-EM data analysis and model building were performed by KE and JB. LH generated preliminary mutational data on alanine variants within the TMDs and supervised GM. Figures were made and edited by GM, ST, KE, JB and MAS. GM and MAS wrote the paper. ST, KE and JB contributed to paper writing.

## ACKNOWLEDGEMENTS

We wish to thank all members of the Seeger and Barandun laboratories for scientific discussions. G.M. and M.A.S. thank the Functional Genomics Center Zurich (Dr. Maria Domenica Moccia, Dr. Lucy Poveda and Dr. Weihong Qi) for their assistance with deep sequencing. We thank S. Štefanić of the Nanobody Service Facility (NSF), University of Zurich for alpaca immunization. Camilo Perez is acknowledged for initial help with grid freezing of EfrCD. The electron microscopy data were collected at the Umeå Core Facility for Electron Microscopy, a node of the Cryo-EM Swedish National Facility, funded by the Knut and Alice Wallenberg Foundation, Erling-Persson Family Foundation, Kempe Foundation, SciLifeLab, Stockholm University, and Umeå University. J.B. acknowledges funding from the Swedish Research council (2019-02011), the SciLifeLab National Fellows program and MIMS. This work was funded by a Swiss National Science Foundation Professorship (PP00P3_144823, to MAS) and a Swiss National Science Foundation project grant (310030_188817, to MAS), an ERC consolidator grant (MycoRailway, no. 772190, to MAS) and an ERC Starting Grant (PolTube, Nr. 948655, to JB).

## Supplementary Information

**Table S1.** Cryo-EM data processing.

|  | **EfrCD:Nb_EfrCD#1** |
| --- | --- |
|  | EMD-12816 |
|  | PDB-7OCY |
| **Data collection and processing** |  |
| Voltage (kV) | 300 |
| Pixel Size (Å) | 1.041 |
| Electron exposure (e-/Å<sup>2</sup>) | 52.1 |
| Defocus range (um) | -1.5 / -3.3 |
| Frames | 40 |
| Symmetry | C1 |
| Initial particle images | 720'893 |
| Final particle images | 212'083 |
| Resolution (Å) | 4.25 |
| FSC threshold | 0.143 |
| Map sharpening B-Factor (Å<sup>2</sup>) | -224.9 |
| **Refinement** |  |
| Initial model used | 4Q4H |
| Model composition |  |
| Non hydrogen Atoms | 7994 |
| Protein residues | 1243 |
| R.m.s deviations |  |
| Bond length (Å) | 0.005 |
| Angles (°) | 1.13 |
| Validation |  |
| MolProbity score | 1.60 |
| Clashscore | 5.26 |
| Poor rotamers (%) | 0.00 |
| Ramachandran |  |
| Favored (%) | 95.37 |
| Allowed (%) | 4.63 |
| Outliers (%) | 0 |

**Table S2.** Characterization of randomly picked depleted cluster variants.

| chain | variant | growth<br>(8 μM<br>daunorubicin) | expression (%)<br>over wt) | Hoechst<br>transport | ethidium<br>transport | ATPase<br>activity in<br>nanodiscs (%)<br>over wt) |
| --- | --- | --- | --- | --- | --- | --- |
| EfrC | F180A | ↓ | 109 |  |  | 251 |
| EfrC | F180W | ↓ | 109 |  |  |  |
| EfrC | F180G | ~wt | 110 |  |  |  |
| EfrC | F180Y | ~wt | 98 |  |  |  |
| EfrC | F180T | ↓ | 121 | <wt | <wt | 152 |
| EfrC | F180K | ↓ | 122 | <wt | <<wt | 104 |
| EfrC | F180C | ↓ | 73 | <wt | <wt |  |
| EfrC | F180M | ~wt | 113 |  |  |  |
| EfrC | N228A | ↓ | 110 |  |  | 170 |
| EfrC | N228S | ↓ | 105 |  |  |  |
| EfrC | N228Y | ↓ | 118 |  |  |  |
| EfrC | N228G | ~wt | 104 |  |  |  |
| EfrC | N228F | ↓ | 125 | ~wt | <wt |  |
| EfrC | N228V | ↓ | 102 | <wt | <wt | 91 |
| EfrC | N228W | ↓ | 128 | <wt | <wt |  |
| EfrC | N228C | ↓ | 100 |  |  |  |
| EfrC | N228E | ↓ | 119 |  |  |  |
| EfrC | N228P | ↓ | 47 |  |  |  |
| EfrC | G232A | ↓ | 121 |  |  | 84 |
| EfrC | G232I | ↓ | 81 | <<wt | <<wt |  |
| EfrC | G232Q | ↓ | 87 | <<wt | <<wt |  |
| EfrC | G232D | ↓ | 97 | <<wt | <<wt | 14 |
| EfrC | G232P | ↓ | 81 |  |  |  |
| EfrC | G232L | ↓ | 88 |  |  |  |
| EfrC | F235A | ↓ | 108 |  |  | 165 |
| EfrC | F235G | ↓ | 107 | <<wt | <<wt |  |
| EfrC | F235Y | ↓ | 53 |  |  |  |
| EfrC | F235W | ~wt | 111 |  |  |  |
| EfrC | F235A | ↓ | 117 | <<wt | <<wt |  |
| EfrC | F235Q | ↓ | 127 | <<wt | <<wt | 137 |
| EfrC | F235H | ↓ | 109 |  |  |  |
| EfrC | F235D | ↓ | 74 |  |  |  |
| EfrC | F235L | ↓ | 104 |  |  |  |
| EfrC | F235E | ↓ | 134 |  |  |  |
| EfrC | F235C | ↓ | 105 |  |  |  |
| EfrD | D136A | ~wt | 34 |  |  | 150 |
| EfrD | D136Y | ↓ | 26 |  |  |  |
| EfrD | D136I | ↓ | 41 | ~wt | ~wt | 106 |
| EfrD | D136T | ~wt | 50 |  |  |  |
| EfrD | D136S | ~wt | 65 |  |  |  |
| EfrD | D136E | ~wt | 97 |  |  |  |
| EfrD | N143A | ↓ | 115 |  |  | 81 |
| EfrD | N143Q | ↓ | 25 |  |  |  |
| EfrD | N143L | ↓ | 1 |  |  |  |
| EfrD | N143S | ↓ | 105 | <wt | <wt | 57 |
| EfrD | N143Y | ↓ | 92 | <wt | <wt |  |
| EfrD | N143R | ↓ | 46 |  |  |  |
| EfrD | N143E | ↓ | 111 | <wt | ~wt |  |
| EfrD | N143A | ↓ | 114 |  |  |  |
| EfrD | N143I | ↓ | 120 |  |  |  |

~wt: wild-type-like growth or transport activity, ↓: decreased growth, <wt : low transport of drug, <<wt : very low transport of drug

**Table S3.**
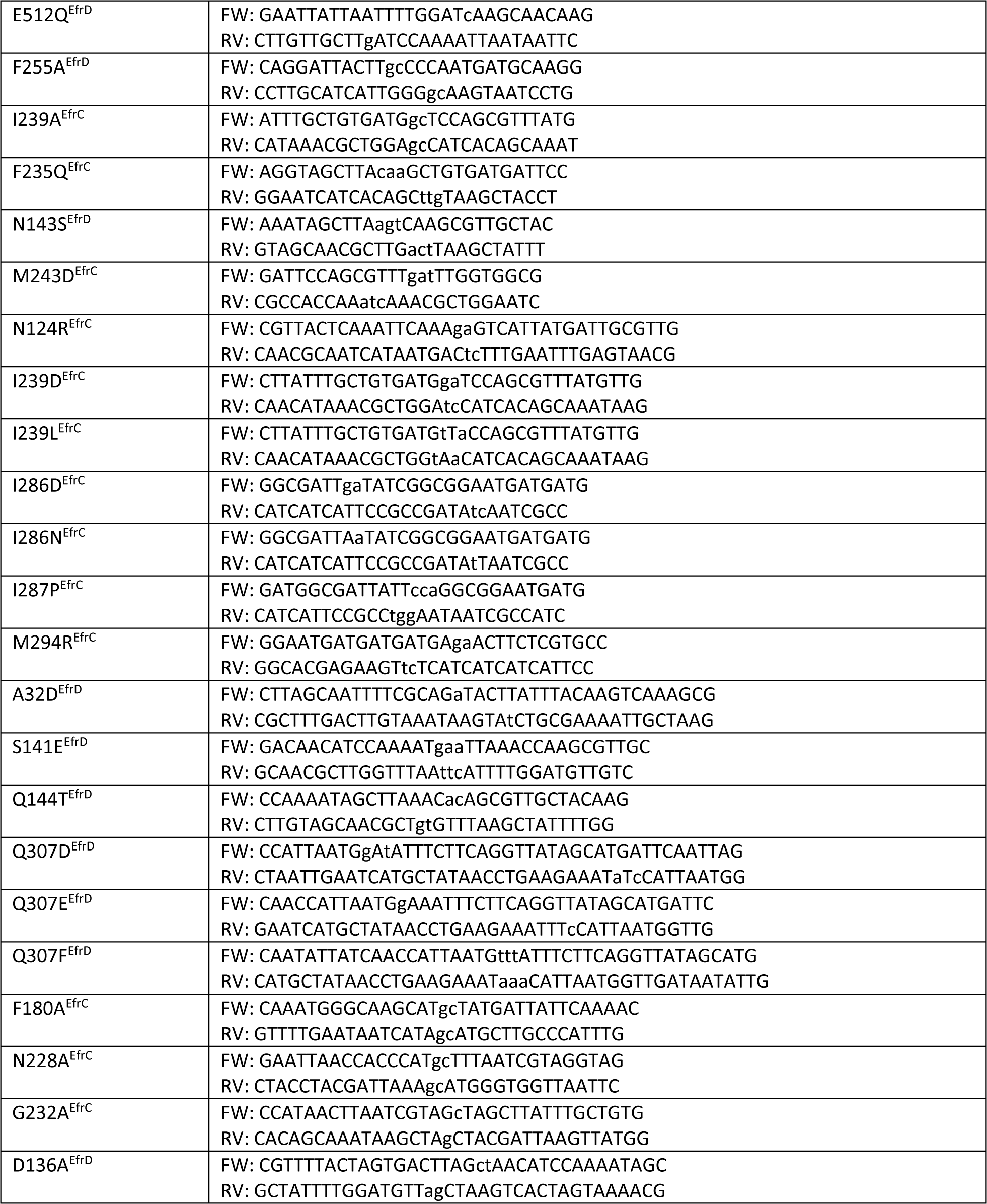
Primer sequences for QuikChange site-directed mutagenesis.

***Supplementary Note 1 – DMS library generation***

To the best of our knowledge, we report here the first successful application of DMS to a drug efflux pump. Therefore, we highlight some technical hurdles, which needed to be overcome in the course of our study.

Under ideal conditions, a DMS input library is evenly distributed. In praxis, however, this is not the case, because every library is to some extent skewed. Consequently, the NGS reads for the 1539 DMS variants of the non-selected input library differ considerably. By increasing the sequencing depth, underrepresented variants can still be read with sufficient read counts, but at increased costs. To achieve an optimal NGS read-out, we made major efforts to render the DMS library as evenly distributed as possible.

In the process of library generation, we encountered and solved the following problems: 1) Technical obstacles in commonly used methods to generate DNA libraries 2) uneven distribution of randomized amino acids 3) diversity bottlenecks and 4) DNA shuffling arising from PCR amplification.

For library generation, our first attempt was to use NNK-randomized primers and overlap PCR to amplify the *efrCD* gene and thereby generate sub-libraries for every randomized position [1]. This approach worked well at the technical level. However, the usage of NNK degenerate primers resulted in skewed, unevenly distributed libraries, because the 20 amino acids are encoded by the 32 NNK codons. Hence, some amino acids (leucine, serine and arginine) are encoded by three codons and are thus overrepresented, whereas the other amino acids are encoded by two or a single codon. Another problem with the NNK-randomization approach is that near-cognate sequencing errors (see Supplementary Note 3) bleed into the reads of randomized codons rather frequently. Therefore, we decided to order the single-site saturation libraries from a commercial provider (Twist Biosciences). These libraries have two technical hallmarks: i) there is only one pre-defined codon for each amino acid (we used codons with highest abundance in *Lactococcus lactis*) and ii) the codons are guaranteed to be fairly evenly distributed for each randomized position (Fig. S4d).

For each of the 81 mutation sites of the *efrCD* gene, we obtained a separate sub-library from Twist Biosciences, which was subcloned one-by-one into the *E. coli* pRexdmsC3GH cloning vector needed as intermediate step for VBEx cloning into a nisin-inducible *Lactococcus lactis* expression vector (Fig. 2a) [2]. In this cloning step, at least 10^3^ colonies were obtained from which the plasmids were prepared, in order to maintain the even distribution of the 19 variant codons for each of the mutation sites.

Next, the DMS library was cloned from the pRexdmsC3GH vector into the *L. lactis* expression vector pNZxDMS (Fig. 2a). The first open reading frame of *efrCD* variants cloned into this vector encodes for (untagged) EfrC, and the second open reading frame encodes for EfrD, followed by 3C protease cleavage site, GFP and a His10-tag. The GFP fusion allowed us to monitor expression levels in cells, whereas the His10-tag and the 3C cleavage site facilitate the purification of individual clones for biochemical characterization. To perform the cloning reaction, pRexdmsC3GH vectors containing the 81 mutation sites were first pooled and then sub-cloned. In a first attempt, we mixed the 81 pRexdmsC3GH sub-libraries at equimolar ratios. However, based on next generation sequencing we realized that the 81 sub-libraries contained variable amounts of remaining wild-type *efrCD* genes, which must originate from library generation by Twist Biosciences using overlap PCR. This resulted in variable frequencies of variant codons for each of the 81 randomized positions (Fig. S4b). To render the library more even in terms of variant codons in each of the 81 randomized positions, we took the NGS reads into account to adjust the mixing ratios of our 81 pRexdmsC3GH sublibraries. As expected, this resulted in a more even distribution of variant codon frequencies of our library (Fig. S4c). In order to measure replicates, we generated three separate DMS pools by mixing three times the respective pREXdmsC3GH vectors (which are referred to as the three biological replicates in this study), followed by three independent vector backbone exchange (VBEx) cloning reactions to finally result in our DMS libraries in the *L. lactis* expression vector pNZdmsC3GH (Fig. 2a, Fig. S4c). To avoid diversity bottlenecks and ensure an even distribution of the randomized codons, the number of colony forming units was at least 500 fold greater than the DMS library size of 1539 possible variants. Since VBEx is solely relying on DNA digestion and ligation and avoids PCR amplification, DNA shuffling and consequential recombination of mutations were avoided [3].

***Supplementary Note 2 – Optimization of selective growth conditions to perform DMS***

The experimental conditions of the selective growth experiment are a critical factor, which required optimization of the drug concentrations and the inducer concentration to adjust the optimal EfrCD expression level. In addition, selection pressures based on cellular growth bear the risk that mutations outside of the plasmid-encoded *efrCD* gene confound the outcome of the DMS experiment. To avoid this, we performed the DMS selections in two growth cycles with a plasmid isolation and retransformation step between the cycles (Fig. 2b).

To achieve optimal competition conditions, we optimized the following parameters: 1) Drug concentrations, 2) expression inducer concentrations and 3) growth conditions. We used a test set of four EfrCD variants, including wild-type EfrCD and inactive E512Q^EfrD^ variant as well as one variant with slightly increased efflux activity (F255A^EfrD^) and one variant with decreased efflux activity (I239A^EfrC^). Of note, these two alanine variants were identified when we performed a preliminary alanine scan of the upper region of the EfrCD drug binding cavity. First, the optimal nisin concentration used to induce *efrCD* expression was determined. A 1:10’000 dilution of a nisin-containing *L. lactis* NZ9700 culture supernatant was found to be optimal, as it leads to high EfrCD overproduction (as measured based on fluorescence of

GFP fused to the EfrD-chain) without affecting cellular growth (Fig. S5a). Further, we identified optimal drug concentrations by performing growth experiments with a test set of variants at various drug concentrations (Fig. S5b). Based on this experiment, we decided to set the following optimal drug concentrations: daunorubicin (8 μM), ethidium (16 μM) and Hoechst (1.5 μM). At these drug concentrations, we enrich for highly active variants (e.g. F255A^EfrD^), partially deplete variants with somewhat decreased efflux activity (e.g. I239A^EfrC^) and fully deplete inactive variants (e.g. E512Q^EfrD^).

When working with plasmid libraries propagated in living organisms, a potential source of artefacts are mutations in the bacterial genome or co-transformation of more than one plasmid per cell, which might provide a cellular growth advantage independently from DMS variants. In addition, by following GFP fluorescence over time, we observed decreased production of EfrCD after prolonged expression (after around 12 generations). To counteract potential sources of artefacts and ensure high EfrCD production levels, our experimental setup includes extraction of plasmid libraries and retransformation into *L. lactis* after one round of growth competition (Fig. 2b).

***Supplementary Note 3 - NGS and data processing pipeline***

NGS turned out to be the limiting (cost) factor of the entire project. A major technical challenge was the length of the *efrCD* genes (around 3.3 kb) and the fact that the mutated residues spread over the entire operon. This prevented us from using Illumina MiSeq (paired end reads of around 250 bp), which is the pre-dominant NGS method used in DMS analyses [4–6]. Instead, we decided to shear the *efrCD* gene into fragments of 130-180 bp length and to sequence them using the Illumina NovaSeq platform as paired-end reads with an overlap of 120-150 bp.

To facilitate future DMS projects, we provide here detailed information on critical aspects of NGS library preparation, next generation sequencing and data analysis (Fig. S6). The input material for NGS-library construction are pNZxDMS plasmids bearing the DMS library before and after competitive growth selection, which are extracted from *L. lactis*. In a first step, these plasmids are digested with SfiI to minimize the amount of backbone plasmid DNA in our NGS analysis. We noted that extraction of the transporter gene without any flanking sequences (using restriction enzymes cutting right at the beginning and end of the *efrCD* gene) resulted in very low sequencing coverage at the 5’-region of *efrC* and the 3’-region of *efrD* (Fig. S4a). By using the SfiI digest, we include around 2x1200 bp of vector backbone upstream and downstream of the *efrCD* gene, which resolved this technical issue. Due to the large size of the *efrCD* transporter genes (3315 bp) we had to shear the DNA to be able to sequence them on Illumina sequencers. An option would be the PacBio technology, which in theory permits for full-length *efrCD* reads [7]. There are techniques, which deal with sequencing of large gene variant libraries such as JigsawSeq, however they rely on heavy randomization of the DNA sequence to generate overlaps to complete the “jigsaw” and are thus not useful for our *efrCD* library with its variant positions being far apart from each other [8]. The latest generation of Illumina next generation sequencers (such as the NovaSeq 6000 system) have Phred Quality Scores of Q30 in more than 90% of bases, corresponding to a base call accuracy of 99.9 %. However, this accuracy was insufficient for our purposes and we decided to rely on overlap paired end sequencing results only (i.e. we only took gene regions into consideration, which were read from both ends with the exact same sequence). For economic reasons, it was more attractive to use the Illumina NovaSeq 6000 system instead of Illumina MiSeq. To enable overlap paired end sequencing, the sheared DNA fragments needed to have a size of around 150 bp, which is considerably shorter than in conventional Illumina library preparations (550 or 350 bp). With some adjustment of the standard Illumina library preparation protocol (see Materials and Methods), we achieved to obtain high quality sequencing libraries with insert sizes of 150 bp.

In our pipeline (Fig. S6a), residual adapter sequences resulting from reading short fragments to the very end are removed. Reads are then aligned to the *efrCD* gene using the burrow wheeler aligner (BWA). Aligned reads are quality filtered by removing reads with low mean quality score (<30) and trimmed to the next codon start. Variants are called from the overlapping sequence of forward and reverse read, thereby only allowing matching reads (i.e. being identical in the forward and reverse read), and sequences with maximally one amino acid substitution with regards to the EfrCD wild-type sequence. The well-established DMS software Enrich2 is then used to calculate enrichment scores by weighted linear least squares regression analysis [9].

Despite considerable progress, sequencing errors of high-throughput sequencers still represent a challenge. Owing to the fact that the paired-end reads are more than 20 times shorter than the entire *efrCD* gene, the great majority of the sequenced fragments are identical to the *efrCD* wild-type sequence and with insufficient accuracy, sequencing errors would be spuriously attributed as mutations. The large number of 81 mutated positions caused an additional technical hurdle. Under idealized circumstances of a perfect DMS input library, NGS in average reads 80 times the wild-type *efrCD* sequence until a mutation is read, which then corresponds to one of the possible 19 substitutions. Hence, NGS reads need to be highly redundant to achieve the required sequencing depth for a statistically significant read-out. In addition, sequencing errors (which cannot be completely suppressed even with overlap paired-end reading) result in near cognate substitutions (exchange of a single base) of the frequently-read wild-type codons, which can “bleed” into the rarely-read codons of true variants.

To quantify the frequency and distribution of sequencing errors for our application, we performed an NGS analysis of the wild-type *efrCD* gene. To this end, we picked a single *L. lactis* colony propagating the *efrCD*-containing plasmid encoding for wild-type EfrCD, checked its correctness with Sanger sequencing and processed it with our DMS pipeline (two independent replicates, prepared and sequenced on two different days). This analysis revealed that only a small subset of our investigated variants (namely 90 out or 1539) were affected by the near-cognate read bleeding issue (Fig. S6b), whereas the remaining 1450 variants exhibited false count rates of 2*10^-5^ or lower. When analyzing the location of the near-cognate read errors by plotting reading errors against the 81 randomized position of the DMS library on the *efrCD* gene, it becomes evident that this issue is highly site-specific (Fig. S6c). Importantly, the data of two independent NGS runs performed on different days with independently prepped wild-type *efrCD* genes are basically identical (Fig. S6c). Finally, the read-errors were analyzed according to all possible base substitutions (Fig. S6d). In line with previous publications [10, 11], we observed C->A and G->T conversions being the most frequently miscalled substitutions.

When we then analyzed our DMS input libraries in light of the near-cognate read bleeding issue, we realized that in around 3 % of the investigated DMS variants, the sequencing errors accounted for > 20 % of the variant reads, which we considered as unacceptably high. Therefore, the corresponding enrichment scores were not calculated and the respective squares are colored in grey in the sequence function maps (Fig. 3).

Finally, we asked the question of how reliable our variant scores were determined. To analyze this, we calculated the variant scores independently for replicate 1 and replicate 2 (see also Fig. 2a how these replicates were generated), and plotted the values against each other (Fig. S6e-g). The resulting pairwise Pearson correlation coefficients were r^2^=0.84 for daunorubicin, r^2^=0.9 for Hoechst and r^2^=0.92 for ethidium, which can be regarded as excellent [9]. Please note that these correlation coefficients were in a similar range when performing the same analysis with replicate 1 versus replicate 3 and replicate 2 versus replicate 3.

---

**Figure S1.**
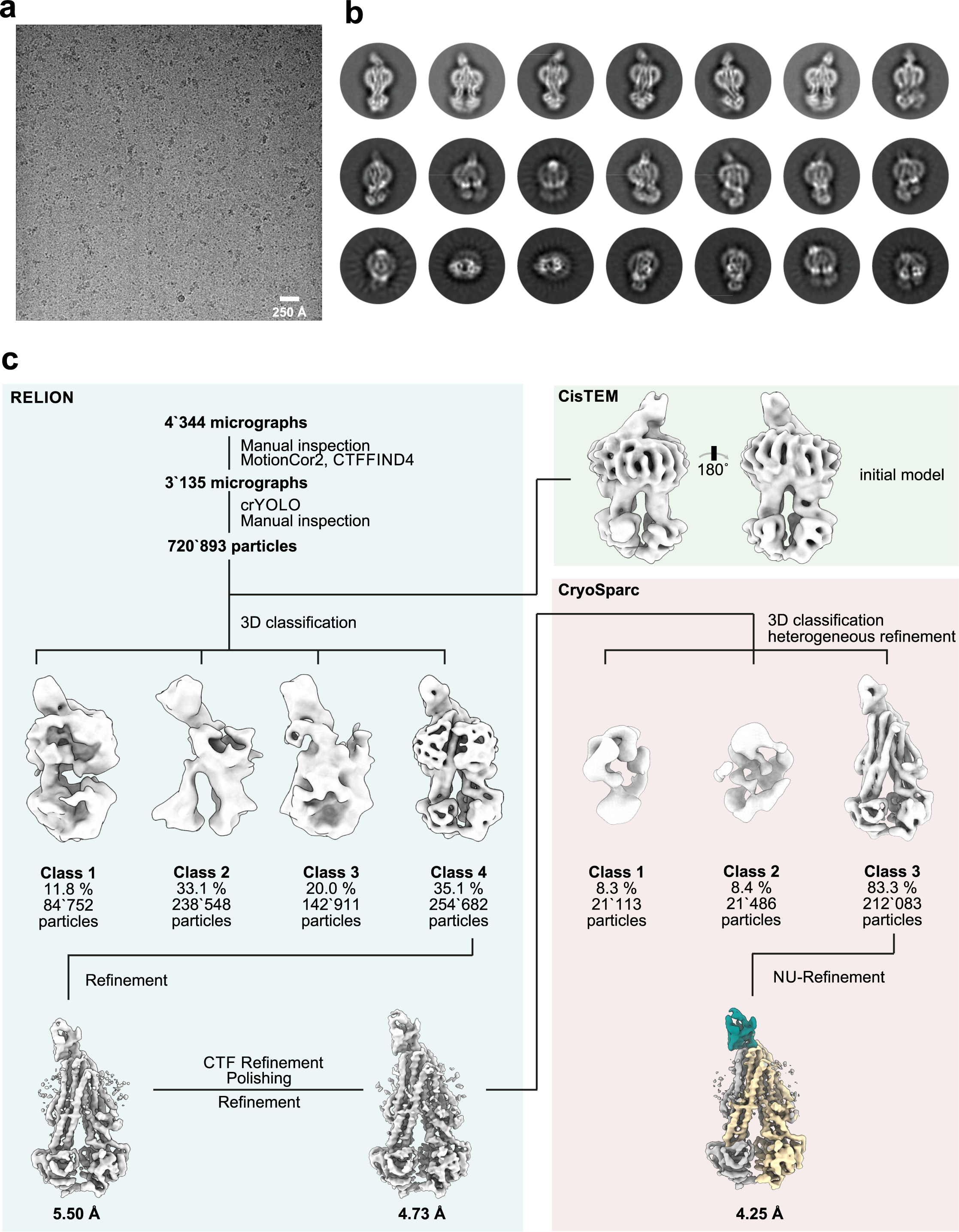
Cryo-electron microscopy data collection and processing. **a,** A representative cryo-EM micrograph of the EfrCD:ATPγS:nanobody sample is shown. The white size bar corresponds to 250 Å. **b,** Selected 2D class averages, calculated using the 254’682 particles from RELION class 4, are displayed. **c,** A schematic classification tree highlights the cryo-EM data processing workflow Intermediate cryo-EM maps are shown in white, while the final 4.25 Å map (EMD-12816) is colored according to individual subunits of the complex. The nanobody is depicted in turquoise, EfrC in silver and EfrD in light yellow. Processing steps performed in RELION have a blue, CisTEM a green and cryoSPARC a pink background.

**Figure S2.**
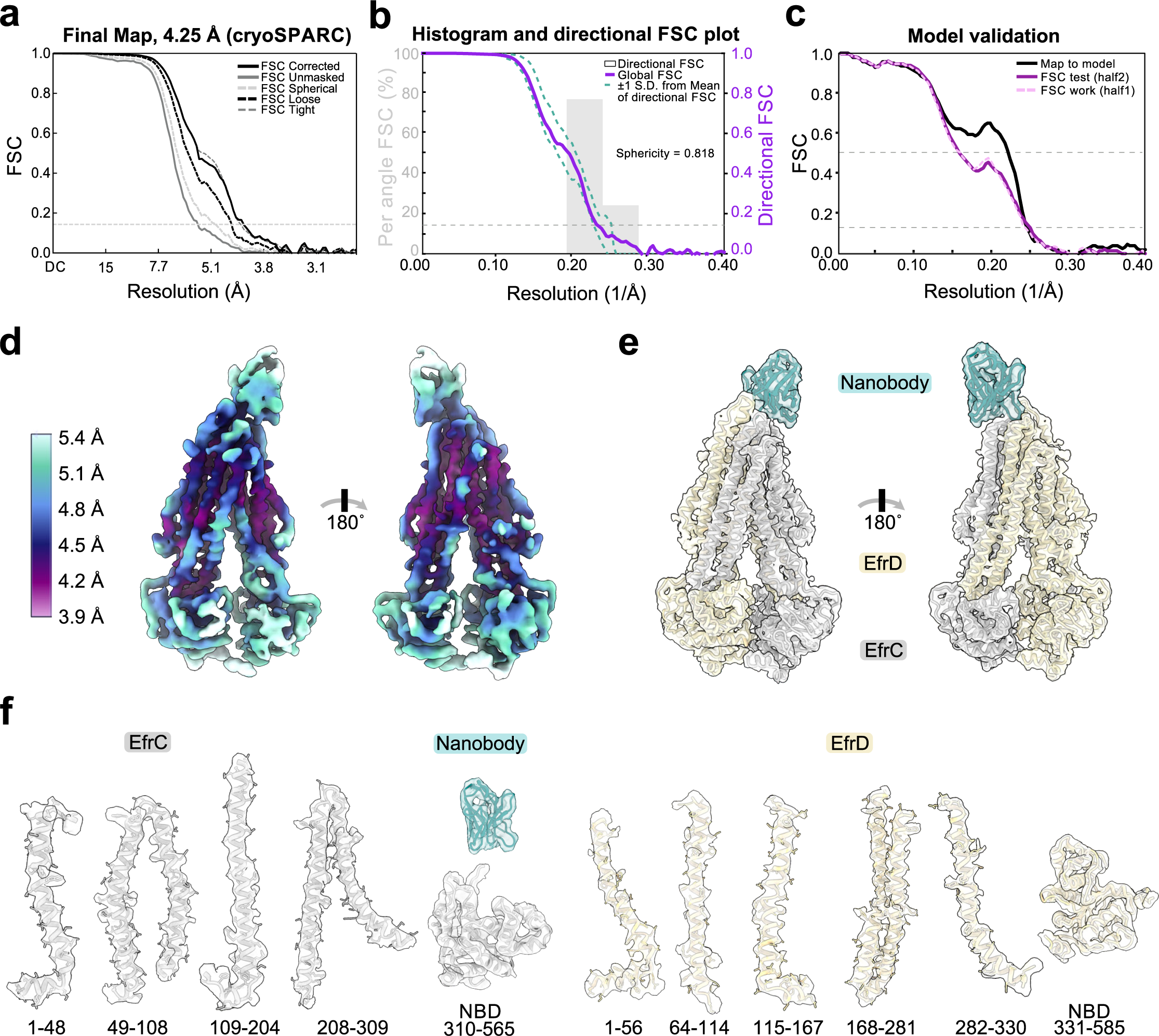
Overall and local resolution estimation, model validation, and cryo-EM density to model fit. **a,** Fourier shell correlation (FSC) curves of the final cryo-EM map (EMD-12816) obtained from cryoSPARC v3.2. **b,** The global half - map FSC (solid purple line) and the spread of the directional resolution values (±1 standard deviation from the mean resolution, teal dotted lines and filled area, right axis) and a histogram of 100 suchvalues, evenly sampled over the 3-dimensional FSC, are shown (grey bars, left axis). A sphericityvalueof 0.816, determined at an FSC threshold of 0.5, suggests isotropic particle distribution. The directional FSC determination was performed using 3DFSC. **c,** Cross-validation of the final refined model and map (black line), and a scrambled model (displacement of all atoms by 0.5 A), refined against half the data, compared against half map 1 (dashed light-purple line) or half map 2 (solid purple line). The thin-dashed line indicates FSC values at 0.143 **(a-c)** or 0.5 **(c). d,** Two 180-degree related views of the final EfrCD reconstruction are shown colored according to local resolution. The local resolution was calculated usingcryoSPARC v3.2. **e,** The overall map-to-model fit between the final refined model (PDB-70CY) and the 4.25 Å cryo-EM map (EMD-12816). The nanobody is colored in turquoise, EfrC in silver, and EfrD in light yellow. **f,** Sections of the cryo-EM map, color-coded according to volume shown in **(e),** are superimposed with the respective molecular models. Subunit names and amino acid number ranges are indicated.

**Figure S3.**
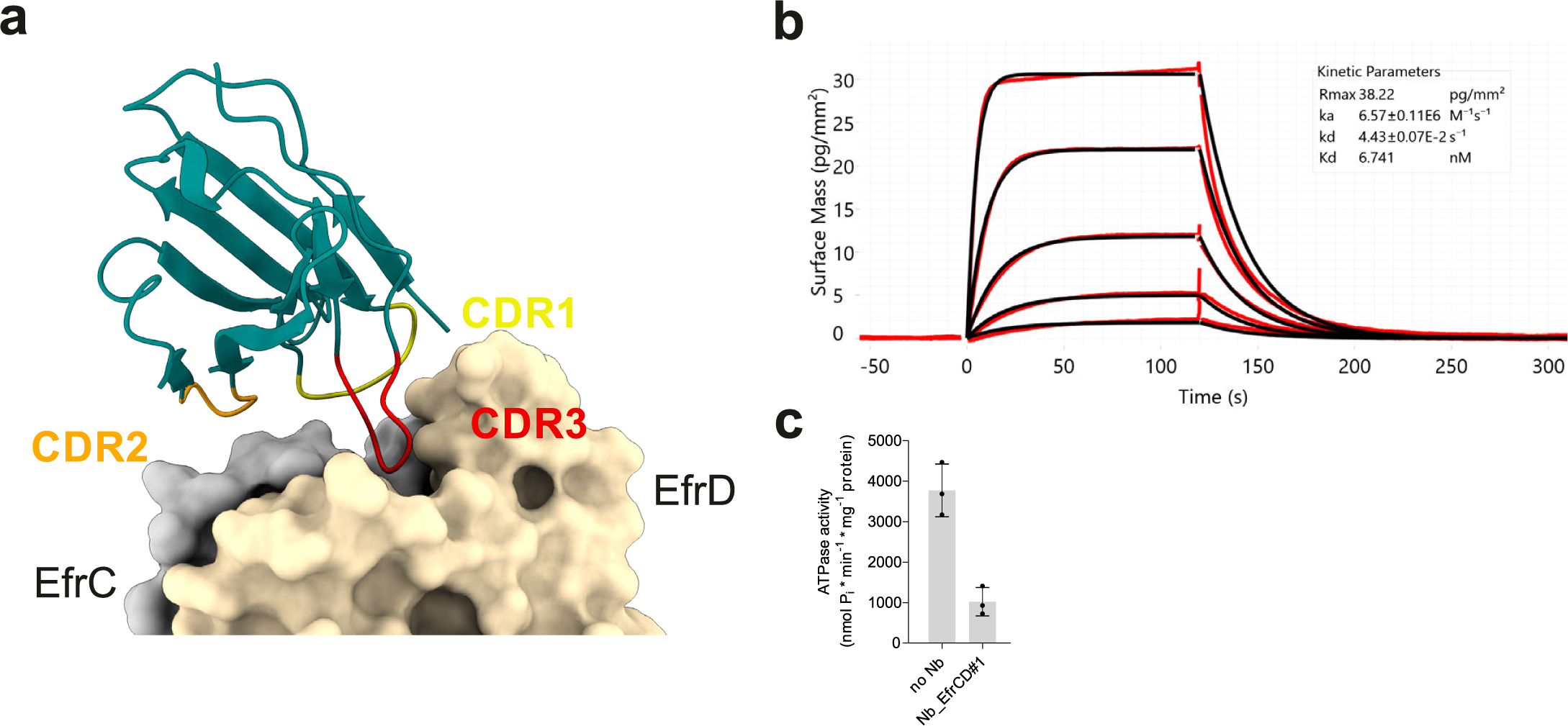
Characterization of nanobody Nb_EfrCD#l. **a,** Cartoon representation of Nb_EfrCD#l binding to the extracellular part of EfrCD. CDRs 1, 2 and 3 are colored in yellow, orange and red, respectively. **b,** Affinity determination of Nb_EfrCD#l usinggrating coupled interferometry (GCI). EfrCD was immobilized on a WAVEchip and Nb_EfrCD#l was injected at 0.33, 1, 3, 9 and 27 nM. A 1:1 kinetic binding model was used for data fitting (black curve) and values for on-rate (*k_a_*), off-rate (*k_d_*) and dissociation constant (*K_D_*) are given in the graph. **c,** ATPase activity of detergent-solubilized EfrCD in the presence and absence of 1 µM Nb_EfrCD#l. Error bars correspond to standard deviations calculated fromtechnical triplicates.

**Figure S4.**
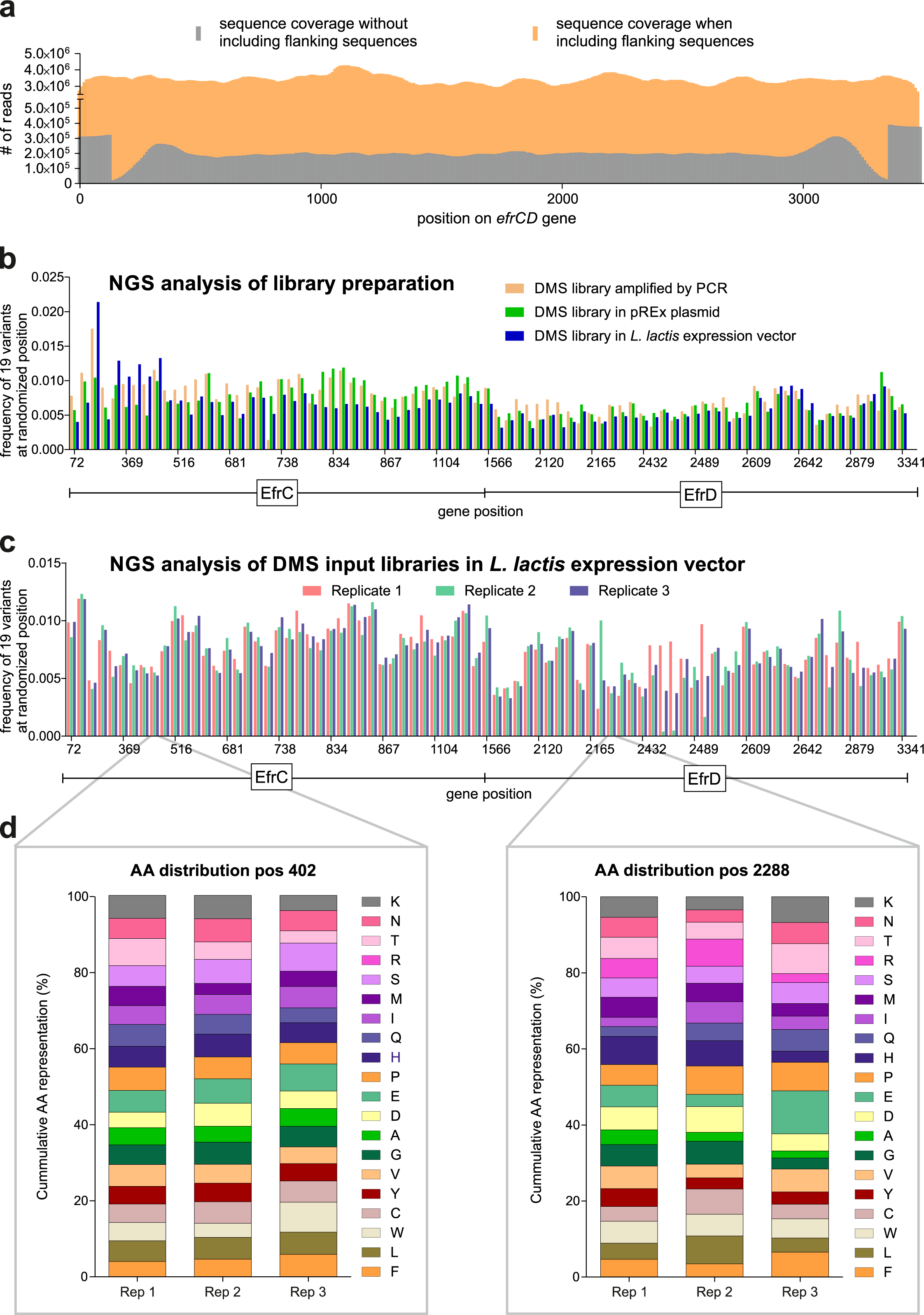
Validation of input libraries. **a,** NGS read coverage of the *efrCD* gene excised without any flanking sequences (grey) versus the inclusion of 2 x 1200 bp flanking sequences up- and downstream of the *efrCD* gene (orange). **b,** Distribution of variant counts for the 81 randomized positions mixed at equimolar ratio. Shown is the NGS analysis of the library as PCR amplified fragments (orange), in the pREx plasmid (green) and in the final *L. lactis* expression vector (blue). **c,** Variant counts of the three input library replicates in the *L. lactis* expression vector used for all experiments. To correct for uneven distribution of variant counts, the ratio of the 81 randomized positions were adjusted based on the variant counts of the NGS analysis shown in **(b). d,** Amino acid distribution at randomized positions 402 (R135^EfrC^) and 2288 (A192^EfrD^) of the *efrCD* gene is shown for the three replicates.

**Figure S5.**
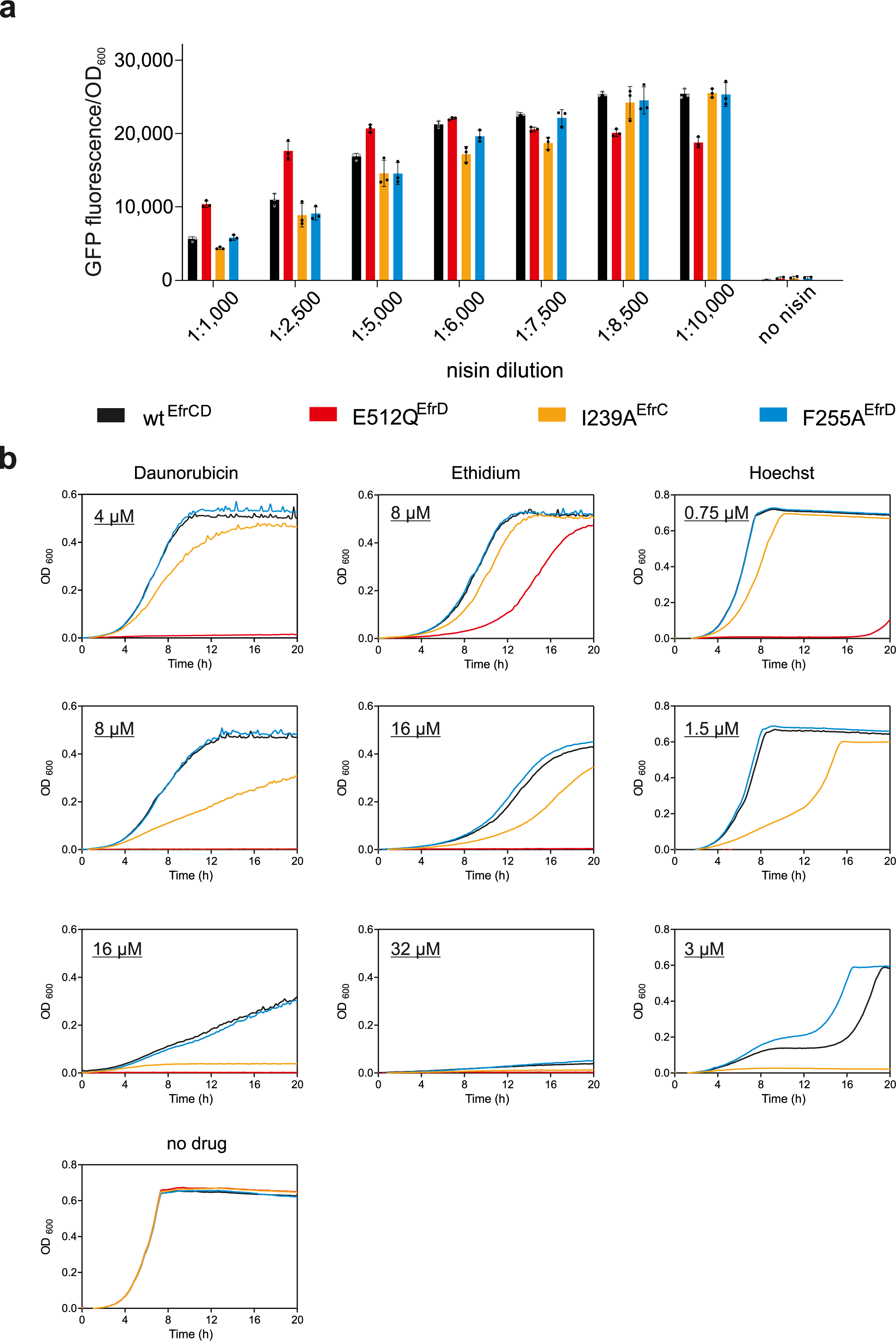
Optimization of competition conditions. **a,** Expression levels assessed by fluorescence of *L. lactis NZ9000ΔlmrCDΔlmrA* expressing EfrCD variants fused to GFP. The test set includes: wild-type EfrCD (black), the inactive E512Q^EfrD^ variant (red), the 1239^EfrC^ variant with decreased (yellow) and the F255A^EFrD^ variant with slightly increased (blue) transport activity compared to wild-type. A nisin­ containing culture supernatant of *L. lactis* NZ9700 was added at varying dilutions. Fluorescence was normalized to the optical density of the cell culture. **b,** Growth of *L. lactis NZ9000ΔlmrCD ΔlmrA* expressing the respective variant upon induction with a 1:10’000 dilution of nisin and in the presence or absence of drugs as indicated. Error bars are standard deviations of technical triplicates.

**Figure S6.**
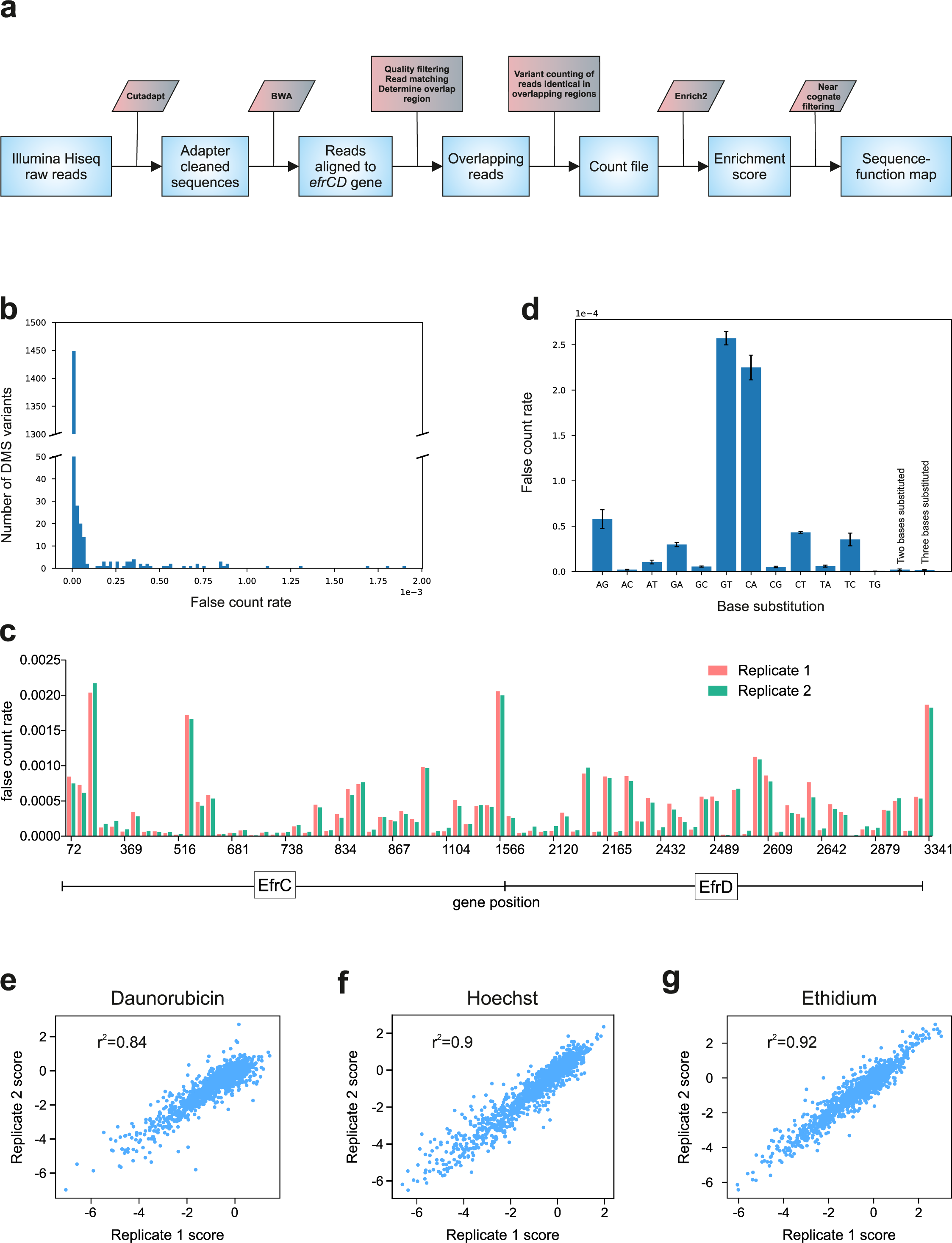
NGS analysis pipeline and validation. **a,** Processes involved in our pipeline are: adapter sequence removal, read alignment using BWA, quality filtering variant counting and enrichment score calculations. **b-d,** Evaluation of sequencing errors using the plasmid encoding for wild-type EfrCD. **b,** Histogram showing the false counts over all 1539 variants of the DMS library. The great majority of variants (around 1450) was not or just slightly affected by sequencing errors, whereas around 90 variants exhibited different levels of sequencing errors. **c,** Error rates plotted versus the 81 randomized positions of the DMS library from two independent sequencing runs of plasmid encoding for wild-type EfrCD (replicate 1 and 2). Sequencing errors are position-specific and highly reproducible. **d,** Error rates of base substitutions. The error bars correspond to the standard deviations for three technical replicates. **e-g,** Pairwise Pearson correlation analysis of variant scores between two replicates selected with daunorubicin **(e),** Hoechst **(fl** and ethidium **(g).** Corresponding correlation analyses with replicate 3 are highly similar (not shown).

**Figure S7.**
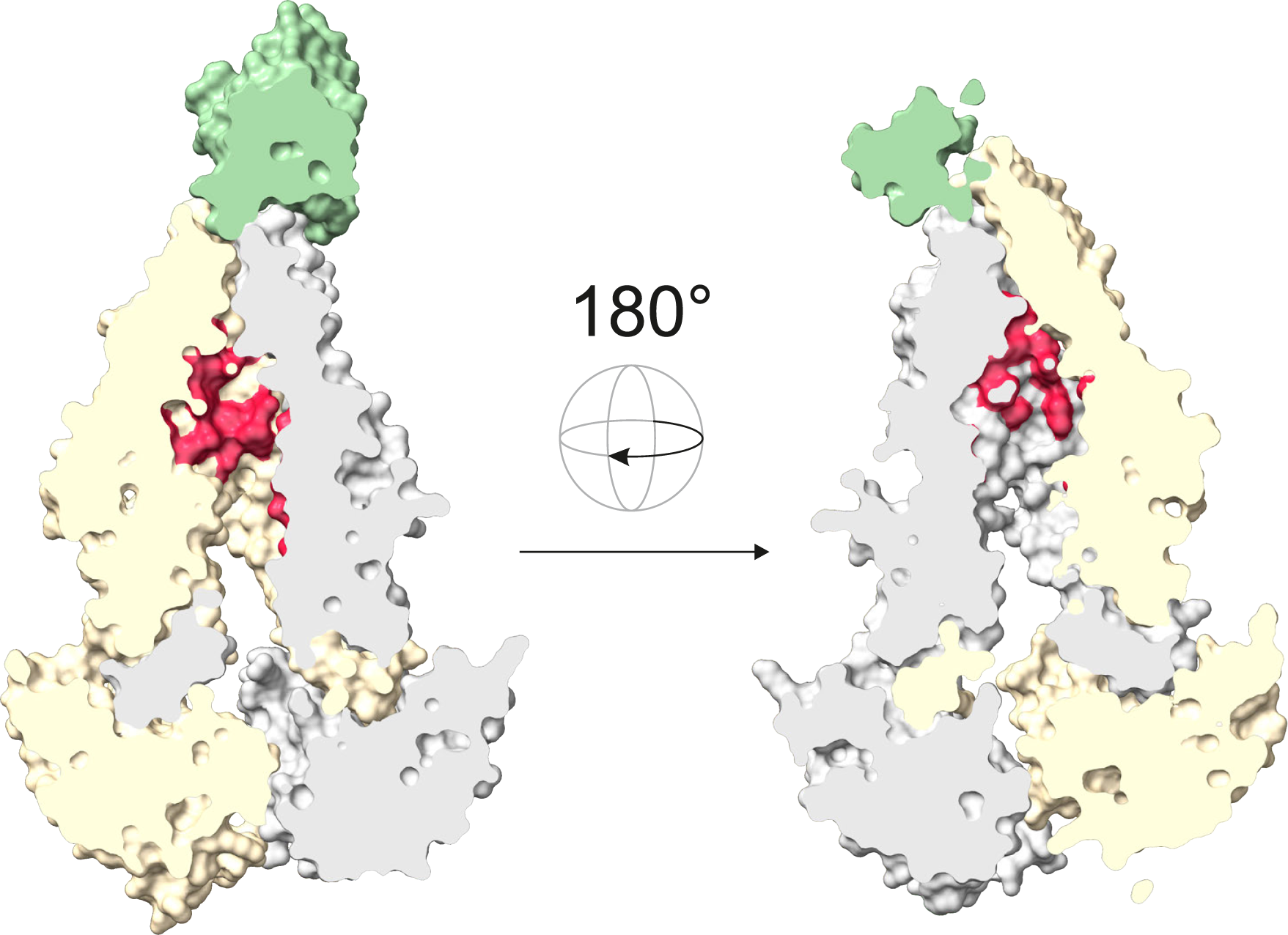
Substitutions towards negatively charged residues enriched in the presence of ethidium. Surface representation of EfrCD cut into two halves to visualize the substrate binding cavity. Residues with enrichment scores > 1 in the presence of ethidium when substituted to aspartate or glutamate are depicted in red.

**Figure S8.**
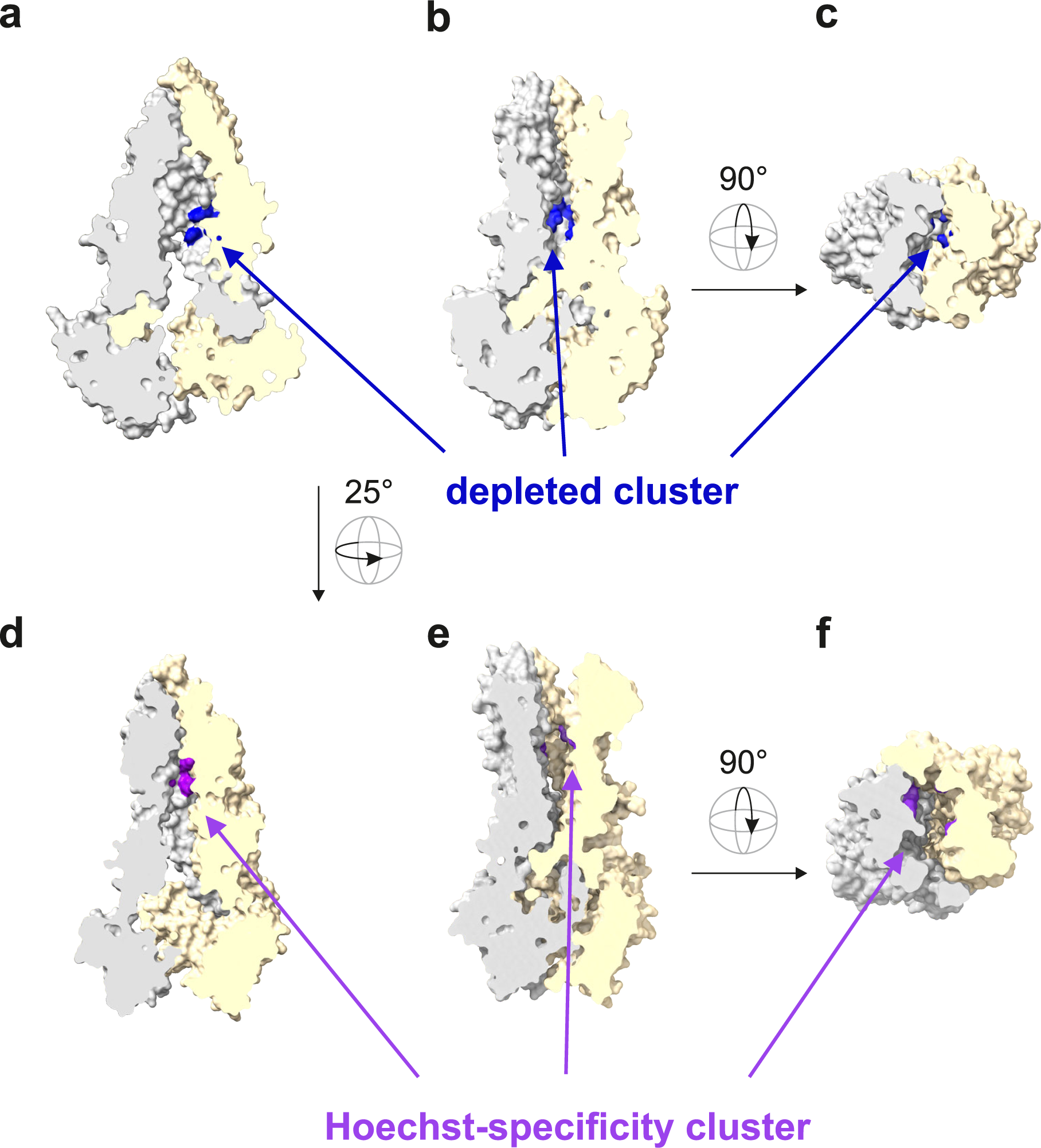
Depleted cluster and Hoechst-specificity cluster in the structural context of outward-facing EfrCD. Side-by-side representation of inward-facing EfrCD structure **(a** and **d)** and outward-facing EfrCD homology model **(b, c, e and f)** based on the coordinates of5av1866 (PDB: 2HYD). The depleted cluster residues are colored in blue **(a-c)** and the Hoechst-specificity cluster residues in purple **(d-f).**

**Figure S9.**
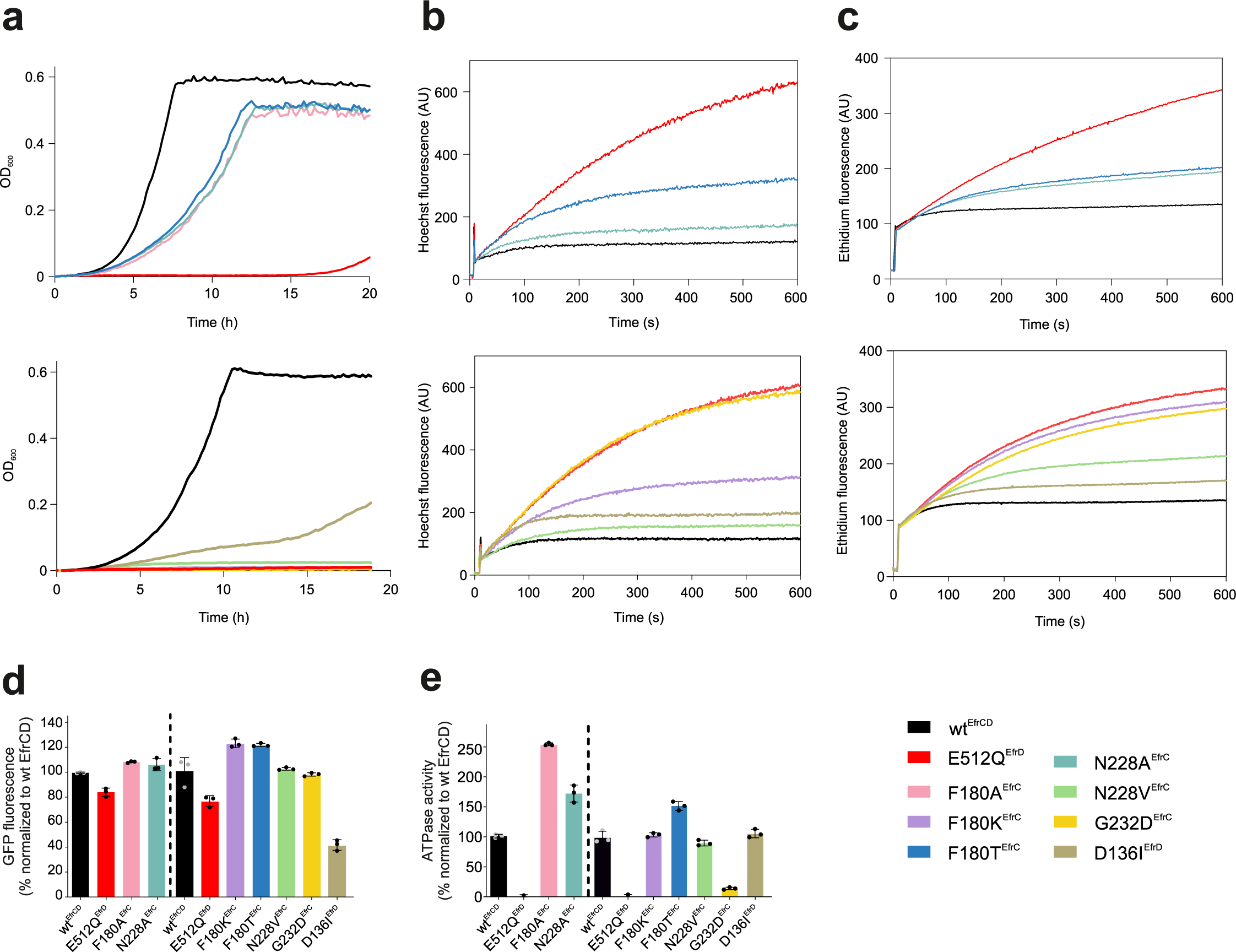
Single clone analysis of depleted cluster variants. **a,** Growth curves of depleted cluster variants determined in the presence of 8 µM daunorubi cin. **b, c,** Transport assay with fluorescent substrates Hoechst **(b)** and ethidium **(c).** Fluorescence spectroscopy was used to monitor the accumulation of drugs by *L. lactis* expressing the respective EfrCD variant. Drugefflux manifests in slower increase of fluorescence. Representative data of two biological replicates are shown. **d,** Protein production level of depleted cluster variants relative to wild-type EfrCD (100 %) asdetermined by GFP-fluorescence in *L. lactis* expressing the variants fused to GFP. e, Basal ATPase activity of purified EfrCD variants measured in nanodiscs. The signal from the inactive E512Q^EfrD^ variant was subtracted and shown is the relative ATPase activity to wild-type EfrCD (100 %). Error bars correspond to standard deviations of technical triplicates. Wild-type EfrCD and the inactive variant E512Q^EfrD^ were included as control in all assays. All fluorescence experiments were carried out twice on the same day (technical duplicates) and were performed on two separate days with freshly prepared cells (biological replicates). Data is shown in two separate graphs for **a-c** and split by a dashed line for **d** and **e** as the variants were analyzed on different days.

**Figure S10.**
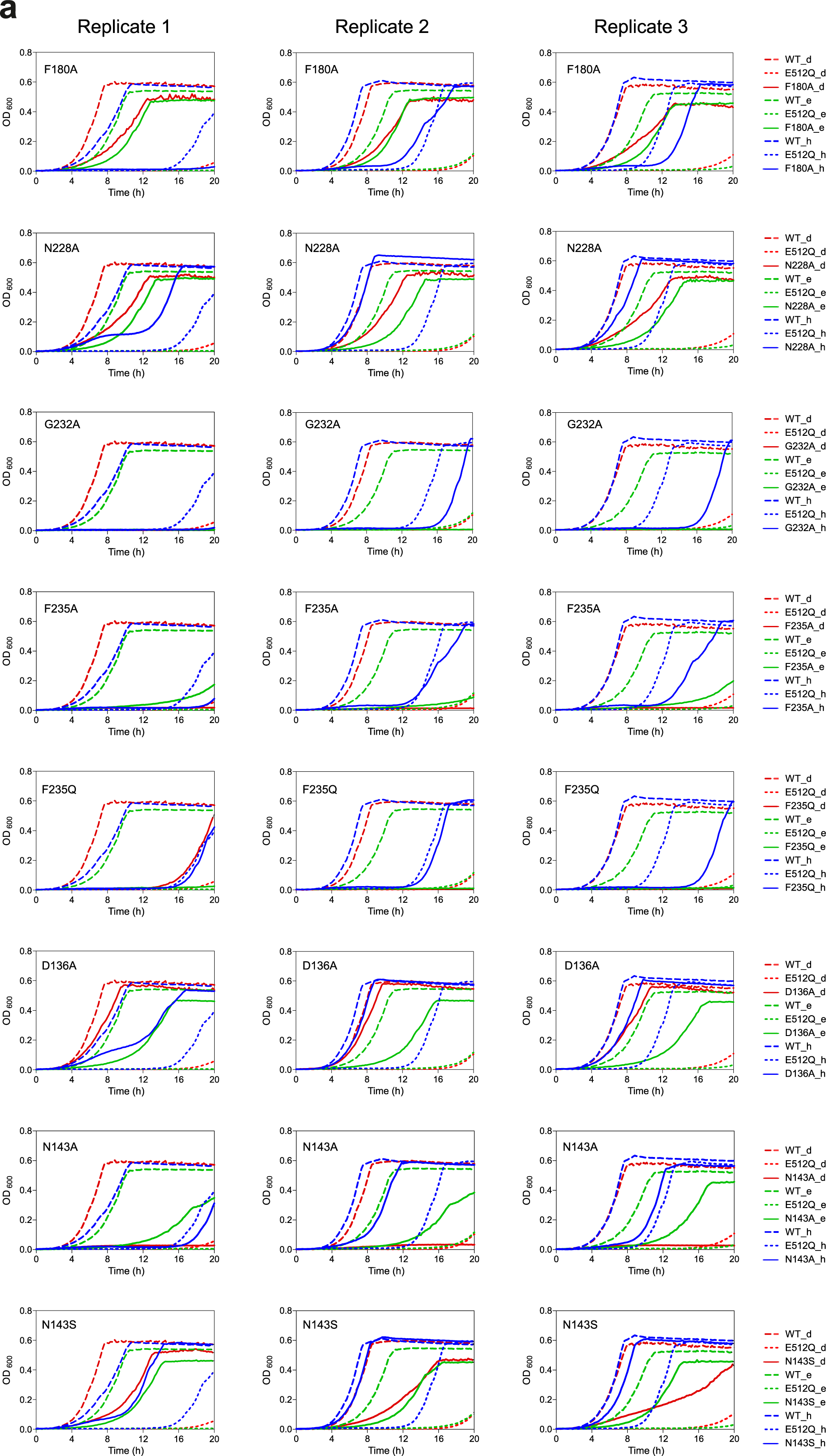
Single clone analysis of depleted cluster variants. **a,** Growth of *L. lactis NZ9000 ΔmrCD ΔmrA* expressing the indicated depleted cluster variants in the presence of daunorubicin (d, red, 8 µM), ethidium (e, green, 16 µM) or Hoechst (h, blue, 1.5 µM) is shown as straight line. Cells expressing wild-type EfrCD (WT, dashed line) or the inactive E512Q^EfrD^ variant (E512Q, dotted line) were included as control. The growth experiment was carried out on three separate days, resulting in three biological replicates.

**Figure S11.**
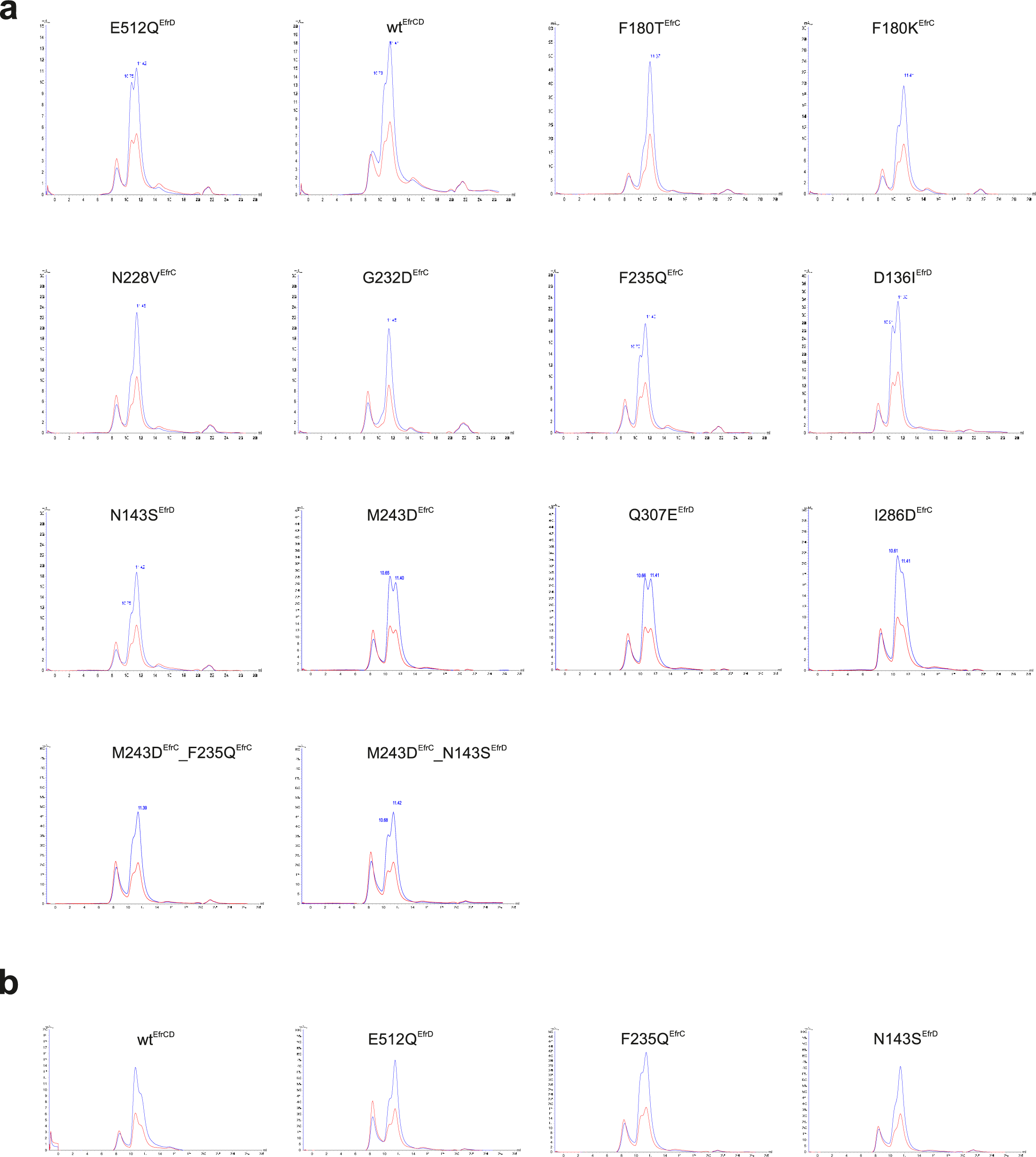
Size exclusion chromatography profiles of detergent-purified EfrCD variants. **a,** EfrCD variants were purified in n-decyl-β-D-maltoside (β-DM) and size exclusion was performed on a Superdex 200 Increase 10/300 GL. Blue and red traces correspond to A_280_ and A_254_, respectively. **b,** Biological replicates of wild-type EfrCD and three variants prepped, and analyzed independently on a different day than in **(a).**

**Figure S12.**
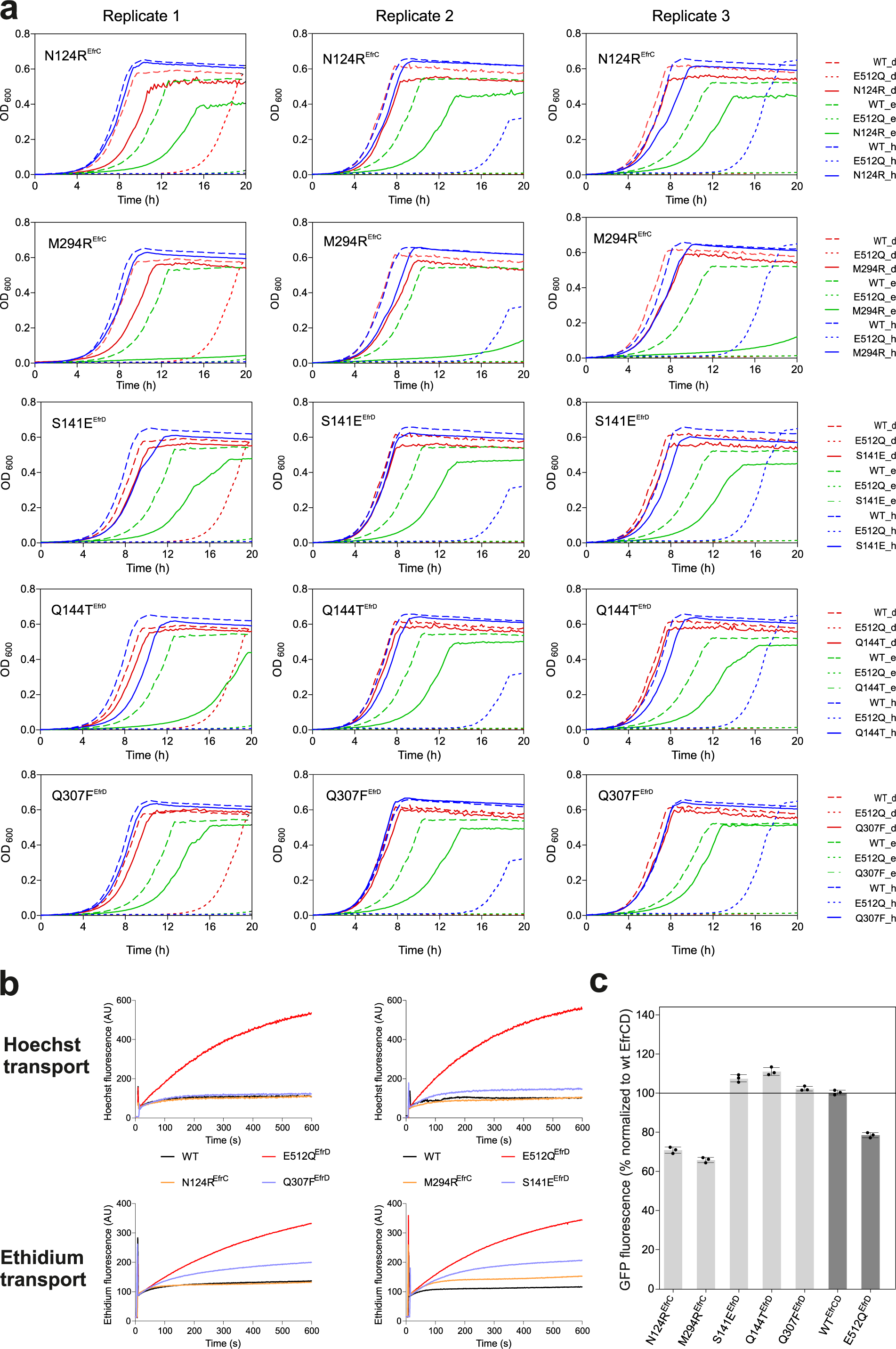
Single clone analysis of ethidium specificity variants. **a,** Growth of *L. lactis NZ9000ΔlmrCD ΔlmrA* expressing the indicated ethidium specificity variants in the presence of daunorubicin (d, red, 8 µM), ethidium (e, green, 16 µM) or Hoechst (h, blue, 1.5 µM) is shown as straight line. Cells expressing wild-type EfrCD (WT, dashed line) or the inactive E512Q^EfrD^ variant (E512Q, dotted line) were included as control. The growth experiment was carried out on three separate days, resulting in three biological replicates. **b,** Accumulation of fluorescent drugs Hoechst 33342 (upper row) or ethidium (lower row) in intact *L. lactis NZ9000 ΔmrCD ΔmrA* expressing ethidium specificity variants. Wild-type EfrCD and inactive E512Q^EfrD^ variant were included in all measurements. Per individual graph, the experimental outcome of the experiments performed on the same day with a batch of equally treated cells is shown. All fluorescence experiments were carried out twice on the same day (technical duplicates) and were performed on two separate days with freshly prepared cells (biological replicates). Representative results from these four replicates are shown. **c,** Expression levels of variants based on GFP-fluorescence of *L. lactis NZ9000 ΔImrCD ΔImrA* expressing transporter variants containing GFP fusion tags. Error bars correspond to standard deviations of technical triplicates.

**Figure S13.**
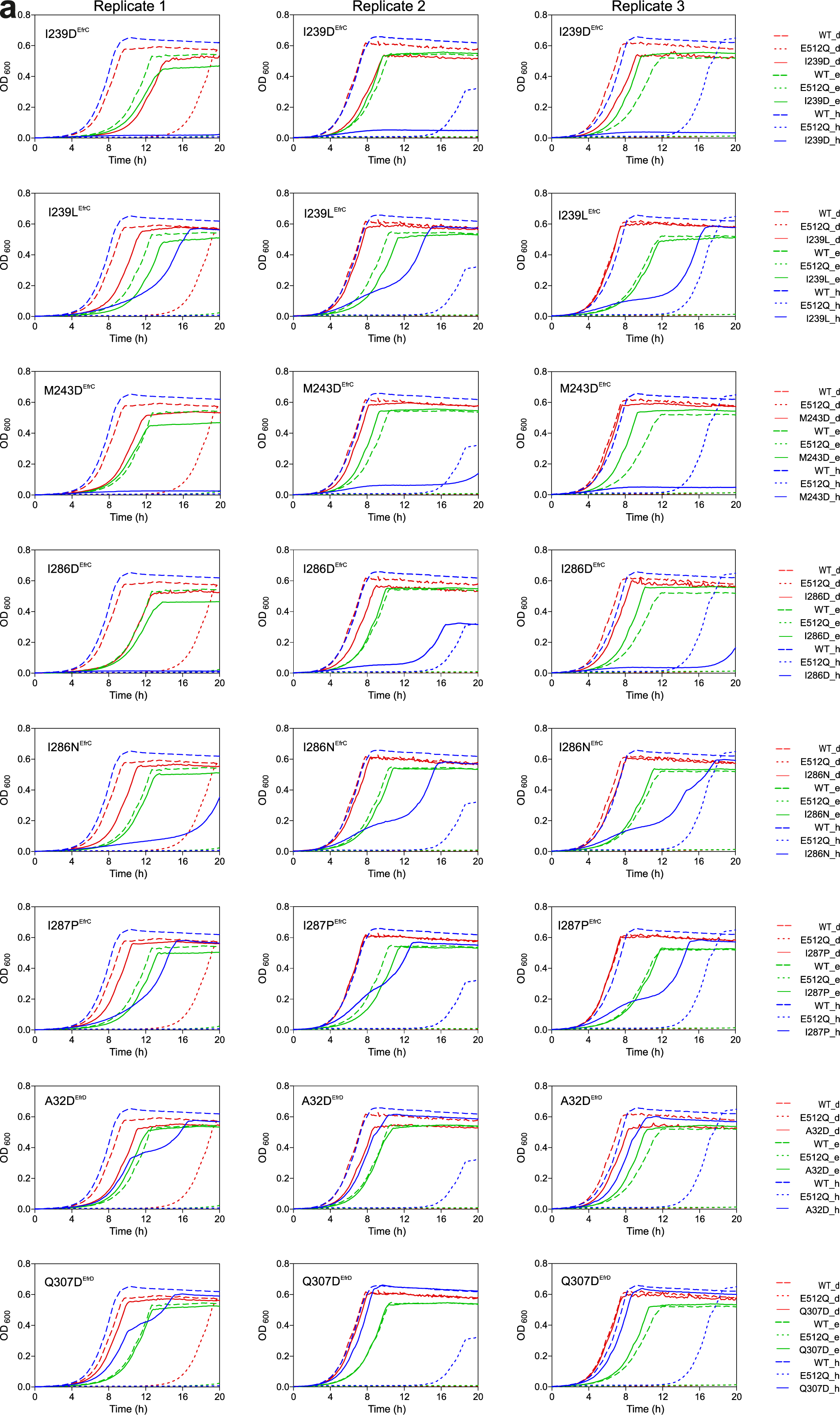
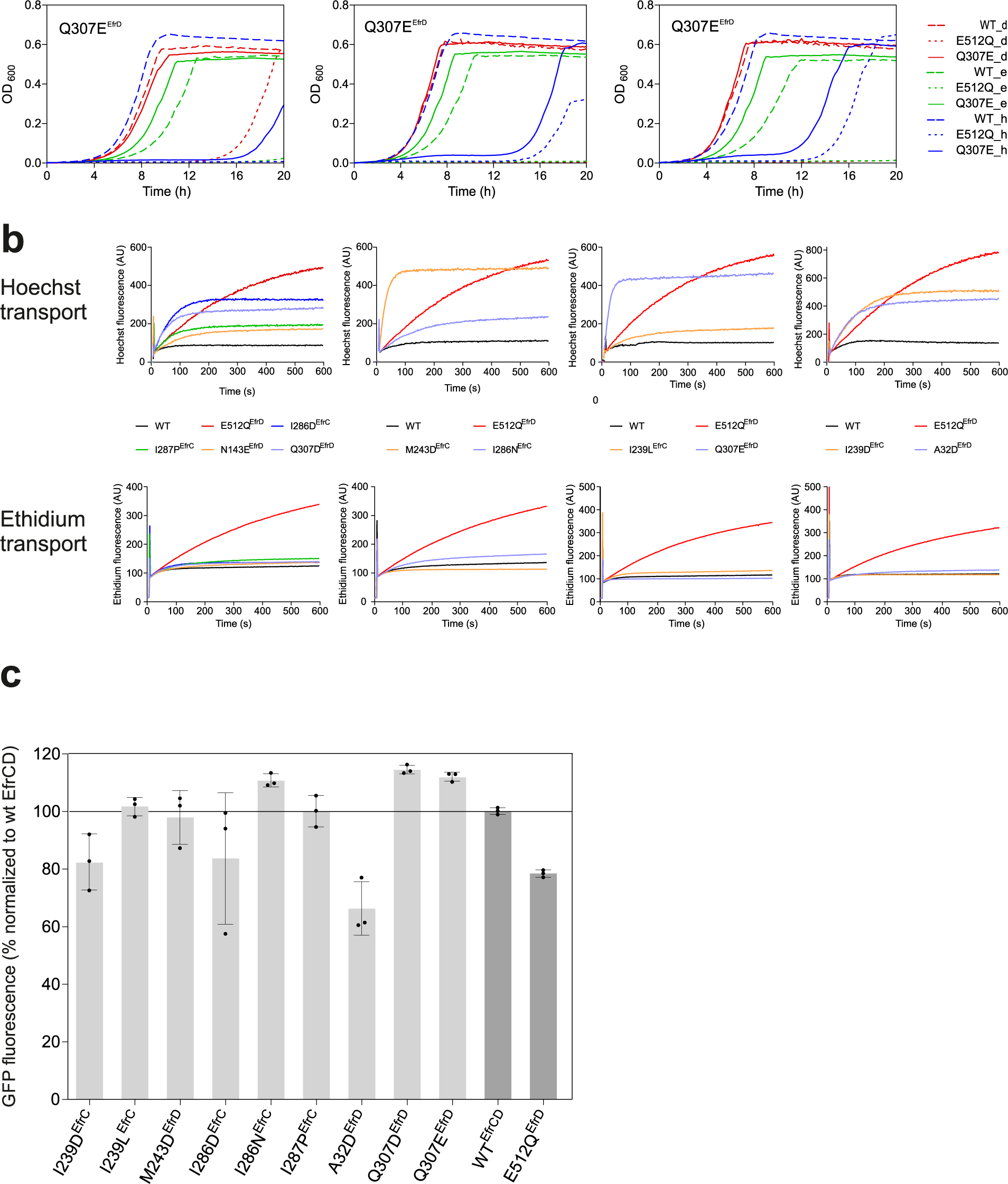
Single clone analysis of Hoechst specificity variants. **a,** Growth of *L. lactis NZ9000 ΔmrCD ΔmrA* expressing the indicated Hoechst specificity variants in the presence of daunorubicin (d, red, 8 µM), ethidium (e, green, 16 µM) or Hoechst (h, blue, 1.5 µM) is shown as straight line. Cells expressing wild-type EfrCD (WT, dashed line) or the inactive E512Q^EfrD^, variant (E512Q, dotted line) were included as control. The growth experiment was carried out on three separate days, resulting in three biological replicates. **b,** Accumulation of fluorescent drugs Hoechst 33342 (upper row) or ethidium (lower row) in intact *L. lactis NZ9000 ΔmrCD ΔmrA* expressing Hoechst specificity variants. Wild-type EfrCD and inactive E512Q^EfrD^ variant were included in all measurements. Per individual graph, the experimental outcome performed on the same day with a batch of equally treated cells is shown. All fluorescence experiments were carried out twice on the same day (technical duplicates) and were performed on two separate days with freshly prepared cells (biological replicates). Representative results from these four replicates are shown. **c,** Expression levels of variants based on GFP-fluorescence of *L. lactis NZ9000 ΔlmrCD ΔlmrA* expressing transporter variants containing GFP fusion tags. Error bars correspond to standard deviations of technical triplicates.

**Figure S14.**
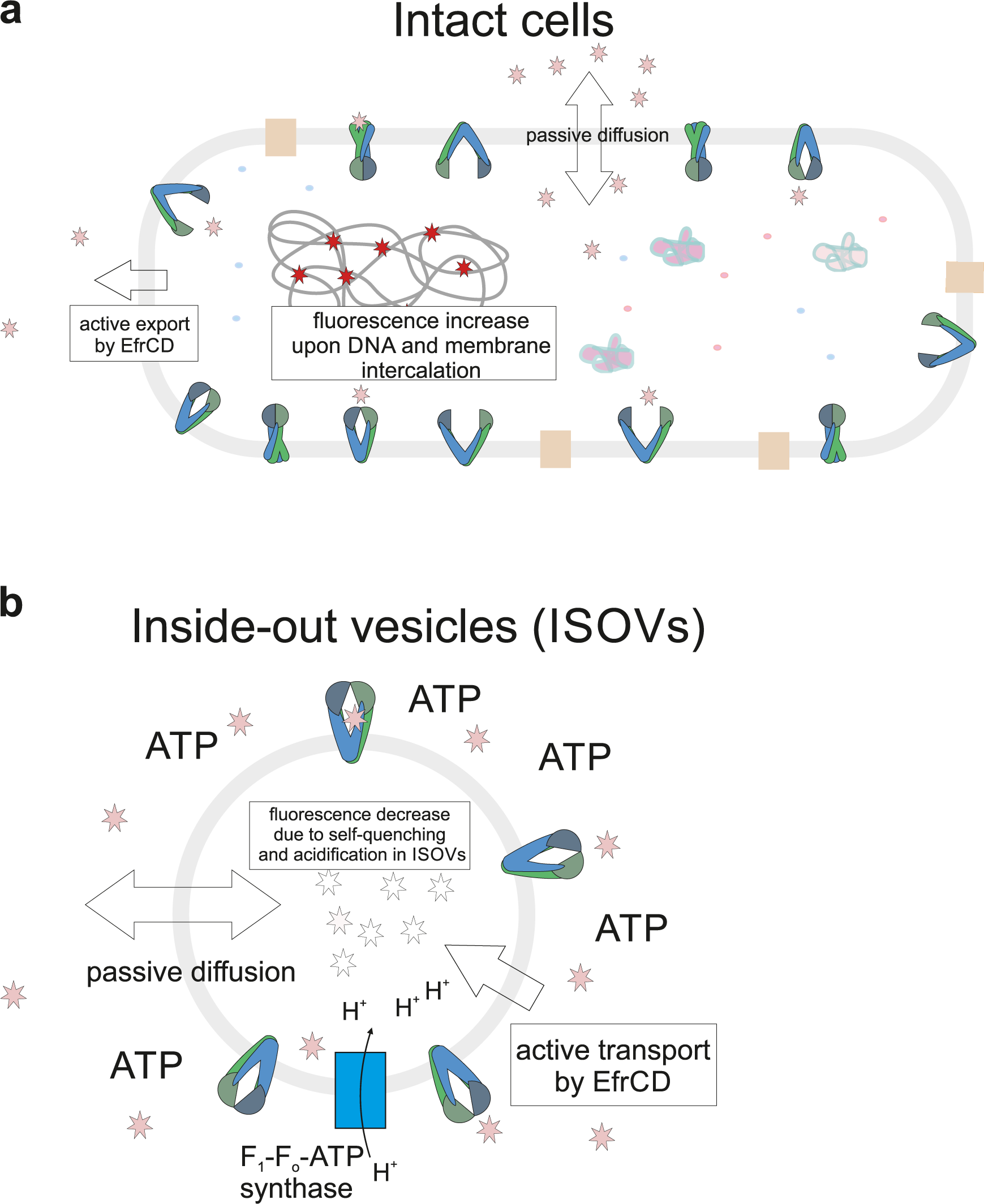
Schematic drawing of Hoechst assays. **a,** Hoechst accumulation assay in intact cells. Hoechst fluorescence increases due to intercalation of Hoechst into thechromosomal DNA and intothelipid bilayer. Active Hoechst efflux mediated by EfrCD results in a slower increase offluorescence. **b,** Hoechst transport into inside-out vesicles (ISOVs). ATP is added from the outside to pump Hoechst into the vesicle lumen. ATP isalso consumed by the F_1_F_0_-ATPaseto pump protons intothe ISOV lumen, thereby acidifyingthe intraluminal milieu. Hoechst fluorescence decreases as a result of self-quenching in the luminal membrane leaflet and protonation of Hoechst due to the lower pHinside the ISOV.

## REFERENCES

1. Theodoulou, F.L. and I.D. Kerr, ABC transporter research: going strong 40 years on. Biochemical Society Transactions, 2015. 43: p. 1033–1040.

2. Du, D., X. Wang-Kan, A. Neuberger, H.W. van Veen, K.M. Pos, L.J.V. Piddock, and B.F. Luisi, Multidrug efflux pumps: structure, function and regulation. Nat Rev Microbiol, 2018. 16(9): p. 523–539.

3. Schumacher, M.A., M.C. Miller, and R.G. Brennan, Structural mechanism of the simultaneous binding of two drugs to a multidrug-binding protein. EMBO J, 2004. 23(15): p. 2923–30.

4. Eicher, T., H.J. Cha, M.A. Seeger, L. Brandstatter, J. El-Delik, J.A. Bohnert, W.V. Kern, F. Verrey, M.G. Grutter, K. Diederichs, and K.M. Pos, Transport of drugs by the multidrug transporter AcrB involves an access and a deep binding pocket that are separated by a switch-loop. Proc Natl Acad Sci U S A, 2012. 109(15): p. 5687–92.

5. Murakami, S., R. Nakashima, E. Yamashita, T. Matsumoto, and A. Yamaguchi, Crystal structures of a multidrug transporter reveal a functionally rotating mechanism. Nature, 2006. 443(7108): p. 173-9.

6. Alam, A., J. Kowal, E. Broude, I. Roninson, and K.P. Locher, Structural insight into substrate and inhibitor discrimination by human P-glycoprotein. Science, 2019. 363(6428): p. 753-+.

7. Le, C.A., D.S. Harvey, and S.G. Aller, Structural definition of polyspecific compensatory ligand recognition by P-glycoprotein. IUCrJ, 2020. 7(4): p. 663–672.

8. Debruycker, V., A. Hutchin, M. Masureel, E. Ficici, C. Martens, P. Legrand, R.A. Stein, H.S. McHaourab, J.D. Faraldo-Gomez, H. Remaut, and C. Govaerts, An embedded lipid in the multidrug transporter LmrP suggests a mechanism for polyspecificity. Nat Struct Mol Biol, 2020. 27(9): p. 829–835.

9. Heng, J., Y. Zhao, M. Liu, Y. Liu, J. Fan, X. Wang, Y. Zhao, and X.C. Zhang, Substrate-bound structure of the E. coli multidrug resistance transporter MdfA. Cell Res, 2015. 25(9): p. 1060–73.

10. Saini, P., T. Prasad, N.A. Gaur, S. Shukla, S. Jha, S.S. Komath, L.A. Khan, Q.M. Haq, and R. Prasad, Alanine scanning of transmembrane helix 11 of Cdr1p ABC antifungal efflux pump of Candida albicans: identification of amino acid residues critical for drug efflux. J Antimicrob Chemother, 2005. 56(1): p. 77–86.

11. Fowler, D.M. and S. Fields, Deep mutational scanning: a new style of protein science. Nat Methods, 2014. 11(8): p. 801–7.

12. Zinkus-Boltz, J., C. DeValk, and B.C. Dickinson, A Phage-Assisted Continuous Selection Approach for Deep Mutational Scanning of Protein-Protein Interactions. Acs Chemical Biology, 2019. 14(12): p. 2757–2767.

13. Romero, P.A., T.M. Tran, and A.R. Abate, Dissecting enzyme function with microfluidic-based deep mutational scanning. Proc Natl Acad Sci U S A, 2015. 112(23): p. 7159–64.

14. Hurlimann, L.M., V. Corradi, M. Hohl, V.G. Bloemberg, D.P. Tieleman, and M.A. Seeger, The Heterodimeric ABC Transporter EfrCD Mediates Multidrug Efflux in Enterococcus faecalis. Antimicrob Agents Chemother, 2016. 60(9): p. 5400–5411.

15. Hürlimann, L.M., M. Hohl, and M.A. Seeger, Split tasks of asymmetric nucleotide-binding sites in the heterodimeric ABC exporter EfrCD. The FEBS Journal, 2017.

16. Hohl, M., L.M. Hurlimann, S. Bohm, J. Schoppe, M.G. Grutter, E. Bordignon, and M.A. Seeger, Structural basis for allosteric cross-talk between the asymmetric nucleotide binding sites of a heterodimeric ABC exporter. Proceedings of the National Academy of Sciences of the United States of America, 2014. 111(30): p. 11025–11030.

17. Rubin, A.F., H. Gelman, N. Lucas, S.M. Bajjalieh, A.T. Papenfuss, T.P. Speed, and D.M. Fowler, A statistical framework for analyzing deep mutational scanning data. Genome Biol, 2017. 18(1): p. 150.

18. Luedtke, N.W., Q. Liu, and Y. Tor, On the electronic structure of ethidium. Chemistry-a European Journal, 2005. 11(2): p. 495–508.

19. Sajid, A., S. Lusvarghi, M. Murakami, E.E. Chufan, B. Abel, M.M. Gottesman, S.R. Durell, and V.S. Ambudkar, Reversing the direction of drug transport mediated by the human multidrug transporter P-glycoprotein. Proceedings of the National Academy of Sciences, 2020. 117(47): p. 202016270.

20. Swain, B.M., D.W. Guo, H. Singh, P.B. Rawlins, M. McAlister, and H.W. van Veen, Complexities of a protonatable substrate in measurements of Hoechst 33342 transport by multidrug transporter LmrP. Scientific Reports, 2020. 10(1).

21. Ambudkar, S.V., I.H. Lelong, J.P. Zhang, C.O. Cardarelli, M.M. Gottesman, and I. Pastan, Partial-Purification and Reconstitution of the Human Multidrug-Resistance Pump - Characterization of the Drug-Stimulatable Atp Hydrolysis. Proceedings of the National Academy of Sciences of the United States of America, 1992. 89(18): p. 8472–8476.

22. Arnold, F.M., M.S. Weber, I. Gonda, M.J. Gallenito, S. Adenau, P. Egloff, I. Zimmermann, C.A.J. Hutter, L.M. Hurlimann, E.E. Peters, J. Piel, G. Meloni, O. Medalia, and M.A. Seeger, The ABC exporter IrtAB imports and reduces mycobacterial siderophores. Nature, 2020. 580(7803): p. 413-+.

23. Al-Shawi, M.K., M.K. Polar, H. Omote, and R.A. Figler, Transition state analysis of the coupling of drug transport to ATP hydrolysis by P-glycoprotein. Journal of Biological Chemistry, 2003. 278(52): p. 52629–52640.

24. Hegedus, C., G. Szakacs, L. Homolya, T.I. Orban, A. Telbisz, M. Jani, and B. Sarkadi, Ins and outs of the ABCG2 multidrug transporter: An update on in vitro functional assays. Advanced Drug Delivery Reviews, 2009. 61(1): p. 47–56.

25. Loo, T.W. and D.M. Clarke, Mutational analysis of ABC proteins. Arch Biochem Biophys, 2008. 476(1): p. 51–64.

26. Tutulan-Cunita, A.C., M. Mikoshi, M. Mizunuma, D. Hirata, and T. Miyakawa, Mutational analysis of the yeast multidrug resistance ABC transporter Pdr5p with altered drug specificity. Genes to Cells, 2005. 10(5): p. 409–420.

27. Srikant, S., R. Gaudet, and A.W. Murray, Selecting for Altered Substrate Specificity Reveals the Evolutionary Flexibility of ATP-Binding Cassette Transporters. Current Biology, 2020. 30(9): p. 1689-+.

28. Schuster, S., S. Kohler, A. Buck, C. Dambacher, A. Konig, J.A. Bohnert, and W.V. Kern, Random Mutagenesis of the Multidrug Transporter AcrB from Escherichia coli for Identification of Putative Target Residues of Efflux Pump Inhibitors. Antimicrobial Agents and Chemotherapy, 2014. 58(11): p. 6870–6878.

29. Swartz, D.J., A. Singh, N. Sok, J.N. Thomas, J. Weber, and I.L. Urbatsch, Replacing the eleven native tryptophans by directed evolution produces an active P-glycoprotein with site-specific, non-conservative substitutions. Scientific Reports, 2020. 10(1).

30. Brown, K., W. Li, and P. Kaur, Role of aromatic and negatively-charged residues of DrrB in multi-substrate specificity conferred by the DrrAB system of Streptomyces peucetius. Biochemistry, 2017: p. acs.biochem.6b01155.

31. Ernst, R., P. Kueppers, C.M. Klein, T. Schwarzmueller, K. Kuchler, and L. Schmitt, A mutation of the H-loop selectively affects rhodamine transport by the yeast multidrug ABC transporter Pdr5. Proc Natl Acad Sci U S A, 2008. 105(13): p. 5069–74.

32. Stockner, T., R. Gradisch, and L. Schmitt, The role of the degenerate nucleotide binding site in type I ABC exporters. FEBS Lett, 2020. 594(23): p. 3815–3838.

33. Mishra, S., B. Verhalen, R.A. Stein, P.C. Wen, E. Tajkhorshid, and H.S. McHaourab, Conformational dynamics of the nucleotide binding domains and the power stroke of a heterodimeric ABC transporter. Elife, 2014. 3: p. e02740.

34. Seeger, M.A., A. Schiefner, T. Eicher, F. Verrey, K. Diederichs, and K.M. Pos, Structural asymmetry of AcrB trimer suggests a peristaltic pump mechanism. Science, 2006. 313(5791): p. 1295-8.

35. Rempel, S., C. Gati, M. Nijland, C. Thangaratnarajah, A. Karyolaimos, J.W. de Gier, A. Guskov, and D.J. Slotboom, A mycobacterial ABC transporter mediates the uptake of hydrophilic compounds. Nature, 2020. 580(7803): p. 409-412.

36. Forsyth, C.M., V. Juan, Y. Akamatsu, R.B. DuBridge, M. Doan, A.V. Ivanov, Z.Y. Ma, D. Polakoff, J. Razo, K. Wilson, and D.B. Powers, Deep mutational scanning of an antibody against epidermal growth factor receptor using mammalian cell display and massively parallel pyrosequencing. Mabs, 2013. 5(4): p. 523–532.

37. Geertsma, E.R. and R. Dutzler, A versatile and efficient high-throughput cloning tool for structural biology. Biochemistry, 2011. 50(15): p. 3272–8.

38. Geertsma, E.R. and B. Poolman, High-throughput cloning and expression in recalcitrant bacteria. Nature methods, 2007. 4(9): p. 705–707.

39. Li, H. and R. Durbin, Fast and accurate short read alignment with Burrows-Wheeler transform. Bioinformatics, 2009. 25(14): p. 1754–1760.

40. Hutter, C.A.J., M.H. Timachi, L.M. Hurlimann, I. Zimmermann, P. Egloff, H. Goddeke, S. Kucher, S. Stefanic, M. Karttunen, L.V. Schafer, E. Bordignon, and M.A. Seeger, The extracellular gate shapes the energy profile of an ABC exporter. Nature Communications, 2019. 10.

41. Zimmermann, I., P. Egloff, C.A.J. Hutter, F.M. Arnold, P. Stohler, N. Bocquet, M.N. Hug, S. Huber, M. Siegrist, L. Hetemann, J. Gera, S. Gmur, P. Spies, D. Gygax, E.R. Geertsma, R.J.P. Dawson, and H.M. Seeger, Synthetic single domain antibodies for the conformational trapping of membrane proteins. Elife, 2018. 7.

42. Zimmermann, I., P. Egloff, C.A.J. Hutter, B.T. Kuhn, P. Brauer, S. Newstead, R.J.P. Dawson, E.R. Geertsma, and M.A. Seeger, Generation of synthetic nanobodies against delicate proteins. Nature Protocols, 2020. 15(5): p. 1707–1741.

43. Zheng, S.Q., E. Palovcak, J.P. Armache, K.A. Verba, Y. Cheng, and D.A. Agard, MotionCor2: anisotropic correction of beam-induced motion for improved cryo-electron microscopy. Nat Methods, 2017. 14(4): p. 331–332.

44. Rohou, A. and N. Grigorieff, CTFFIND4: Fast and accurate defocus estimation from electron micrographs. J Struct Biol, 2015. 192(2): p. 216–21.

45. Wagner, T., F. Merino, M. Stabrin, T. Moriya, C. Antoni, A. Apelbaum, P. Hagel, O. Sitsel, T. Raisch, D. Prumbaum, D. Quentin, D. Roderer, S. Tacke, B. Siebolds, E. Schubert, T.R. Shaikh, P. Lill, C. Gatsogiannis, and S. Raunser, SPHIRE-crYOLO is a fast and accurate fully automated particle picker for cryo-EM. Commun Biol, 2019. 2: p. 218.

46. Grant, T., A. Rohou, and N. Grigorieff, cisTEM, user-friendly software for single-particle image processing. Elife, 2018. 7.

47. Zivanov, J., T. Nakane, B.O. Forsberg, D. Kimanius, W.J. Hagen, E. Lindahl, and S.H. Scheres, New tools for automated high-resolution cryo-EM structure determination in RELION-3. Elife, 2018. 7.

48. Punjani, A., J.L. Rubinstein, D.J. Fleet, and M.A. Brubaker, cryoSPARC: algorithms for rapid unsupervised cryo-EM structure determination. Nat Methods, 2017. 14(3): p. 290–296.

49. Punjani, A., H. Zhang, and D.J. Fleet, Non-uniform refinement: adaptive regularization improves single-particle cryo-EM reconstruction. Nat Methods, 2020. 17(12): p. 1214–1221.

50. Adams, P.D., P.V. Afonine, G. Bunkoczi, V.B. Chen, N. Echols, J.J. Headd, L.W. Hung, S. Jain, G.J. Kapral, R.W. Grosse Kunstleve, A.J. McCoy, N.W. Moriarty, R.D. Oeffner, R.J. Read, D.C. Richardson, J.S. Richardson, T.C. Terwilliger, and P.H. Zwart, The Phenix software for automated determination of macromolecular structures. Methods, 2011. 55(1): p. 94–106.

51. Tan, Y.Z., P.R. Baldwin, J.H. Davis, J.R. Williamson, C.S. Potter, B. Carragher, and D. Lyumkis, Addressing preferred specimen orientation in single-particle cryo-EM through tilting. Nature Methods, 2017. 14(8): p. 793-+.

52. Brown, A., F. Long, R.A. Nicholls, J. Toots, P. Emsley, and G. Murshudov, Tools for macromolecular model building and refinement into electron cryo-microscopy reconstructions. Acta Crystallographica Section D-Structural Biology, 2015. 71: p. 136–153.

53. Tang, G., L. Peng, P.R. Baldwin, D.S. Mann, W. Jiang, I. Rees, and S.J. Ludtke, EMAN2: An extensible image processing suite for electron microscopy. Journal of Structural Biology, 2007. 157(1): p. 38–46.

54. Schrodinger, LLC, The PyMOL Molecular Graphics System. 2015.

55. Goddard, T.D., C.C. Huang, E.C. Meng, E.F. Pettersen, G.S. Couch, J.H. Morris, and T.E. Ferrin, UCSF ChimeraX: Meeting modern challenges in visualization and analysis. Protein Sci, 2018. 27(1): p. 14–25.

## REFERENCES

1. Kille, S., C.G. Acevedo-Rocha, L.P. Parra, Z.G. Zhang, D.J. Opperman, M.T. Reetz, and J.P. Acevedo, Reducing codon redundancy and screening effort of combinatorial protein libraries created by saturation mutagenesis. ACS Synthetic Biology, 2013. 2(2): p. 83–92.

2. Geertsma, E.R. and B. Poolman, High-throughput cloning and expression in recalcitrant bacteria. Nature methods, 2007. 4(9): p. 705–707.

3. Egloff, P., I. Zimmermann, F.M. Arnold, C.A.J. Hutter, D. Morger, L. Opitz, L. Poveda, H.A. Keserue, C. Panse, B. Roschitzki, and M.A. Seeger, Engineered peptide barcodes for in-depth analyses of binding protein libraries. Nature Methods, 2019. 16(5): p. 421–428.

4. Starita, L.M. and S. Fields, Deep Mutational Scanning: Library Construction, Functional Selection, and High-Throughput Sequencing. Cold Spring Harb Protoc, 2015. 2015(8): p. 777–80.

5. Newberry, R.W., J.T. Leong, E.D. Chow, M. Kampmann, and W.F. DeGrado, Deep mutational scanning reveals the structural basis for alpha-synuclein activity. Nat Chem Biol, 2020. 16(6): p. 653–659.

6. Jones, E.M., N.B. Lubock, A.J. Venkatakrishnan, J. Wang, A.M. Tseng, J.M. Paggi, N.R. Latorraca, D. Cancilla, M. Satyadi, J.E. Davis, M.M. Babu, R.O. Dror, and S. Kosuri, Structural and functional characterization of G protein-coupled receptors with deep mutational scanning. Elife, 2020. 9.

7. Rhoads, A. and K.F. Au, *PacBio Sequencing and Its Applications.* Genomics Proteomics Bioinformatics, 2015. 13(5): p. 278–89.

8. Cho, N., B. Hwang, J.K. Yoon, S. Park, J. Lee, H.N. Seo, J. Lee, S. Huh, J. Chung, and D. Bang, De novo assembly and next-generation sequencing to analyse full-length gene variants from codon-barcoded libraries. Nat Commun, 2015. 6: p. 8351.

9. Rubin, A.F., H. Gelman, N. Lucas, S.M. Bajjalieh, A.T. Papenfuss, T.P. Speed, and D.M. Fowler, A statistical framework for analyzing deep mutational scanning data. Genome Biol, 2017. 18(1): p. 150.

10. Pfeiffer, F., C. Gröber, M. Blank, K. Händler, M. Beyer, J.L. Schultze, and G. Mayer, Systematic evaluation of error rates and causes in short samples in next-generation sequencing. Scientific Reports, 2018. 8(1): p. 1–14.

11. Ma, X., Y. Shao, L. Tian, D.A. Flasch, H.L. Mulder, M.N. Edmonson, Y. Liu, X. Chen, S. Newman, J. Nakitandwe, Y. Li, B. Li, S. Shen, Z. Wang, S. Shurtleff, L.L. Robison, S. Levy, J. Easton, and J. Zhang, Analysis of error profiles in deep next-generation sequencing data. Genome Biology, 2019. 20(1): p. 50.

